# The Y14-p53 Regulatory Circuit in Megakaryocyte Differentiation and Thrombocytopenia

**DOI:** 10.1101/2020.11.03.367508

**Authors:** Chun-Hao Su, Wei-Ju Liao, Wei-Chi Ke, Ruey-Bing Yang, Woan-Yuh Tarn

## Abstract

Thrombocytopenia-absent radius syndrome is caused by a deletion in chromosome 1q21.1 in *trans* with *RBM8A* mutations in the noncoding regions. We generated megakaryocyte-specific *Rbm8a* knockout (*Rbm8a*KO^MK^) mice that exhibited marked thrombocytopenia, internal hemorrhage, and splenomegaly, indicating a disorder of platelet production. *Rbm8a*KO^MK^ mice accumulated immature megakaryocytes in the bone marrow and spleen. Depletion of Y14/RBM8A in human erythroleukemia (HEL) cells inhibited phorbol ester-induced polyploidy and downregulated the signaling pathways associated with megakaryocyte maturation. Accordingly, *Rbm8a*KO^MK^ mice had reduced expression of surface glycoproteins on platelets and impaired coagulation. Moreover, p53 level was increased in Y14-depleted HEL cells and *Rbm8a*KO^MK^ megakaryocytes. Treatment with a p53 inhibitor restored *ex vivo* differentiation of *Rbm8a*KO^MK^ megakaryocytes and unexpectedly activated Y14 expression in HEL cells. Knockout of *Trp53* in part restored the platelet count of *Rbm8a*KO^MK^ mice. These results indicate that the Y14-p53 circuit plays a critical role in megakaryocyte differentiation and platelet production.

## INTRODUCTION

Thrombocytopenia-absent radii (TAR) syndrome is a rare congenital disorder characterized by bilateral radial aplasia and a reduced platelet count (Albers et al., 2013). An early study has revealed that thrombocytopenia of the TAR syndrome results from defective differentiation of megakaryocyte precursors (Letestu et al., 2000). Genetic studies have then indicated that TAR is caused by heterozygosity for a deletion within chromosome 1q21.1 (encompassing *RBM8A*) in *trans* with mutations in regions involved in regulating *RBM8A* (Albers et al., 2012). The deletion and mutations together result in diminished *RBM8A* expression in platelets from TAR syndrome patients (Albers et al., 2013), suggesting that *RBM8A* deficiency impacts bone morphogenesis and platelet production.

*RBM8A* encodes the RNA processing factor Y14/RBM8A (hereafter referred to as Y14) that participates in multiple steps of mRNA metabolism. Y14 is a constituent of the exon junction complex, which marks the position of the splice junction in mature mRNAs and initiates mRNA surveillance during the pioneer round of translation (Maquat et al., 2010). Depletion of Y14, however, alters mRNA splicing patterns, indicating that Y14 also modulates alternative splicing (Fukumura et al., 2016; Michelle et al., 2012). Moreover, Y14 is involved in DNA damage repair through its direct interaction with the non-homologous end joining machinery (Chuang et al., 2019). Y14 deficiency leads to accumulation of DNA damage and cell-cycle arrest at G2/M phase (Lu et al., 2017). *Rbm8a* haploinsufficiency impairs mouse cortical development due to reduced numbers of progenitors and neurons and the perturbation of cortical lamination (Mao et al., 2015). Morpholino-mediated knockdown of *rbm8a* in *Danio rerio* results in disorganized myofibers and motor axon outgrowth defects (Gangras et al., 2020). To date, however, the impact, if any, of *RBM8A* deficiency on platelet abundance remains unknown.

Platelets arise from cytoplasmic blebbing and fragmentation of mature megakaryocytes. Megakaryopoiesis is a multi-step process consisting of the commitment of hematopoietic stem cells to the megakaryocytic lineage, megakaryocyte maturation, and terminal differentiation (Machlus and Italiano Jr., 2013; Mazzi et al., 2018). Promegakaryoblasts undergo endomitosis to increase their DNA content ranging from 2N to 128N (64N in mice). The endomitotic process is an abortive mitosis without telophase and cytokinesis, giving rise to a polylobulated nucleus. A number of cell-cycle regulators have been implicated in endomitosis, such as the cyclin-dependent kinase inhibitors p21^CIP1/WAF1^ and p19^INK4D^ and the anaphase-promoting complex regulator Cdc20 (Raslova et al., 2007; Taniguchi et al., 1999; Trakala et al., 2015). Ablation of *Cdc20* in megakaryocytes through Pf4-Cre (Pf4, platelet factor 4) reduces ploidy level and impairs platelet production in mice (Trakala et al., 2015). The tumor suppressor p53 controls both cell-cycle progression and apoptosis. In contrast to *Cdc20* knockout, however, *TP53*-knockout megakaryocytes exhibit increased ploidy and differentiation potential *in vitro* (Apostolidis et al., 2012a). Overexpression or pharmacological activation of p53 leads to decreased polyploidization of megakaryocytes (Apostolidis et al., 2012b; Mahfoudhi et al., 2016). Therefore, Cdc20 and p53 play opposite roles in polyploidization of megakaryocytes. In light of Y14 deficiency-induced p53 activation (Lu et al., 2017; Mao et al., 2016), we postulated that Y14 has a role in megakaryocyte differentiation, which may involve p53.

First, to explore the role of Y14 in megakaryocytic differentiation and platelet production, we generated megakaryocyte-specific *Rbm8a* knockout mice and knocked down Y14 expression in an erythroleukemia cell line. *Rbm8a* knockout mice indeed exhibited thrombocytopenia. We then evaluated whether knockout or inhibition of p53 could reverse the effect of Y14 deficiency on megakaryocyte differentiation.

## RESULTS

### Megakaryocyte-Specific *Rbm8a* Knockout Causes Thrombocytopenia

To ablate *Rbm8a* in the megakaryocyte lineage, we generated a *loxP*-flanked *Rbm8a* allele using a CRISPR/Cas9-mediated insertion system (Figure 1A). Wild-type, heterozygous and homozygous floxed alleles were distinguished by PCR (Figure 1B). Mice carrying homologous *loxP*-flanked *Rbm8a* alleles were mated to Pf4-Cre mice (Con^+^); the offspring were born in an approximately Mendelian ratio as assessed 2 weeks after birth (Figure 1C, and Table S1). Because Pf4-Cre drives megakaryocyte and platelet-restricted knockout, we examined Y14 expression in sections of bone marrow and spleen from 9-week-old heterozygous and homozygous knockouts, *i.e*., Pf4^Cre/+^;*Rbm8a*^f/+^ (Con) and Pf4^Cre/+^;*Rbm8a*^f/f^ (*Rbm8a*KO^MK^), respectively. Immunohistochemical (IHC) staining revealed a significant reduction of Y14 protein in *Rbm8a*KO^MK^ megakaryocytes as compared with control samples (Figure 1D), indicating efficient depletion of Y14 in megakaryocytes.

**Figure 1.**
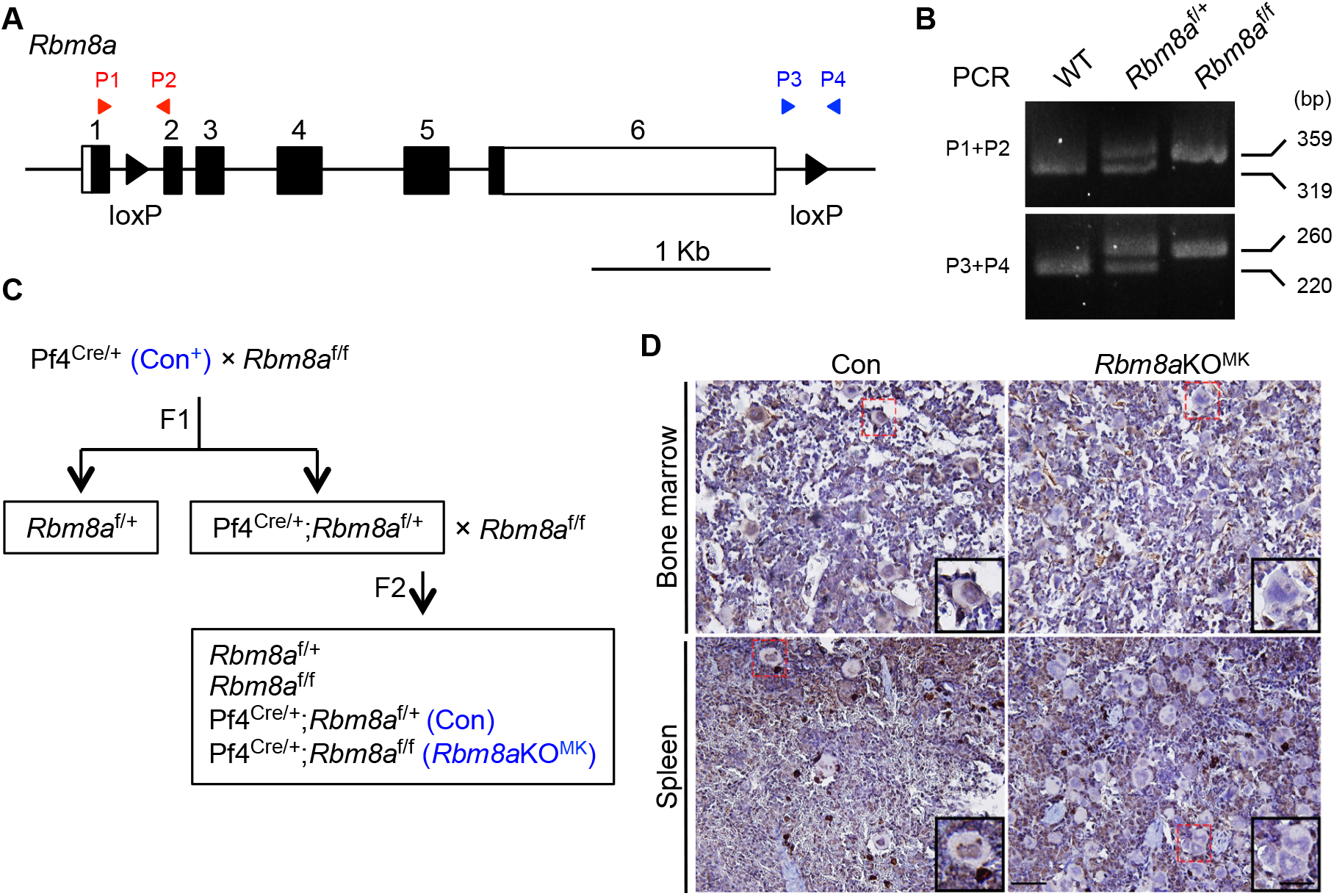
Generation of Megakaryocyte-Specific *Rbm8a* Knockout Mice. (A) Schematic diagram of the mouse *Rbm8a* gene containing six exons (black boxes, coding regions; white boxes, untranslated regions); black arrowheads in intron 1 and the 3’ untranslated region represent *loxP* sites that were generated with the CRISPR/Cas9 system. (B) Genotyping of *Rbm8a*-floxed mice using the indicated primer pairs (red and blue arrowheads in panel A). WT, wild type. (C) Schematic diagram illustrating the generation of megakaryocyte-specific *Rbm8a* knockout mice. Pf4^Cre/+^ (Pf4-Cre, Con^+^); Pf4^Cre/+^;*Rbm8a*^f/+^ (Con); Pf4^Cre/+^;*Rbm8a*^f/f^ (*Rbm8a*KO^MK^). (D) Immunohistochemical staining for Y14 in sections of bone marrow and spleen from 9-week-old Con (Pf4^Cre/+^;*Rbm8a*^f/+^) and *Rbm8a*KO^MK^ mice. A selected megakaryocyte in the red dashed-line square is magnified in the inset. Scale bars represent 50 μm (insets 25 μm).

Some of the male *Rbm8a*KO^MK^ mice died within 2-3 weeks after birth; the death rate was 90% at 8 weeks of age (Figure 2A). Such a high mortality rate may have resulted from subcutaneous and gastrointestinal bleeding that was frequently observed in 2-week-old male mice, indicating platelet defects and perhaps inflammation (Figure 2B); still, other possibilities cannot be excluded. Although all female *Rbm8a*KO^MK^ mice survived until the end of the study (up to 3 months), they showed moderate subdermal hemorrhage and internal hemorrhage near lymph nodes at 8 weeks of age (Figure 2C). Moreover, *Rbm8a*KO^MK^ mice of both genders exhibited splenomegaly (Figure 2D), which may have resulted from sequestration of immature megakaryocytes in the spleen (see below). We also observed a drastic reduction in the number of platelets in *Rbm8a*KO^MK^ mice at 6-8 weeks of age, including one male (Figure 2E). Although it has been argued that PF4 may be minimally expressed in hematopoietic stem cells as well as lymphoid and myeloid cells (Gollomp and Poncz, 2019), the count of all other types of blood cells remained normal in *Rbm8a*KO^MK^ mice (Figure 2E and Table S2). *Rbm8a*KO^MK^ mice showed a robust increase in tail bleeding compared with control mice (Figure 2F). A severe reduction in platelet number likely accounted for both the prolonged tail bleeding time and internal bleeding of *Rbm8a*KO^MK^ mice. This result indicated an important role for *Rbm8a* in platelet production.

**Figure 2.**
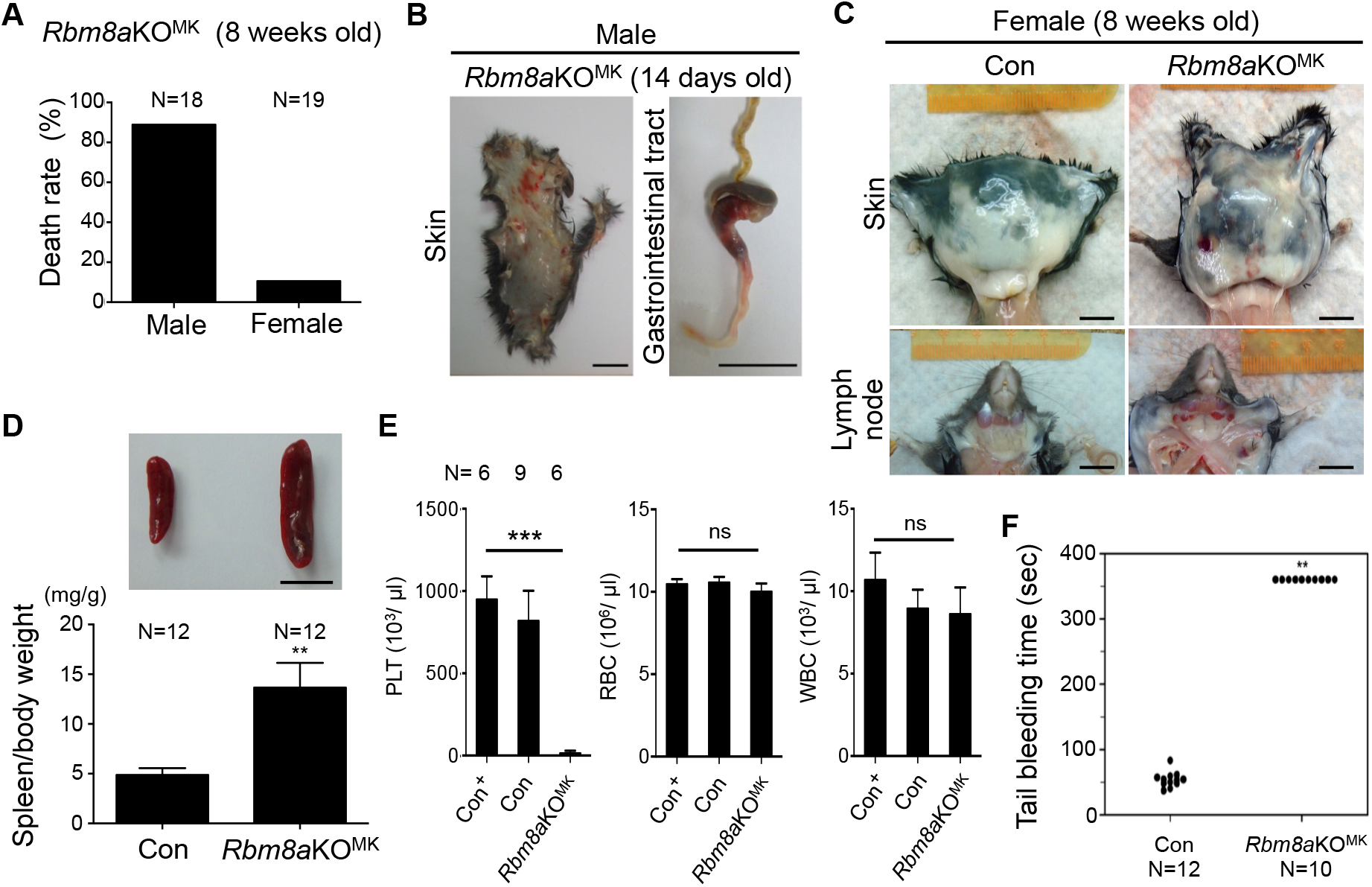
Megakaryocyte-Specific *Rbm8a* Knockout Causes a Reduction in Platelet Count and Related Pathological Phenotypes. (A) Death rate of 8-week-old male and female *Rbm8a*KO^MK^ mice. (B) Subcutaneous tissue (skin) and gastrointestinal tract of 14-day-old male *Rbm8a*KO^MK^mice. (C) Subcutaneous tissue and lymph nodes of 8-week-old female Con and *Rbm8a*KO^MK^ mice. (D) The spleen of 10- to 12-week-old mice as in panel C. Bar graph shows the spleen/body weight ratio (mg/g; mean ± SEM, N=12 in each group, **p < 0.01). Scale bars in panels B, C, and D represent 1 cm. (E) Hematological analysis of blood from 6- to 8-week-old mice of the indicated genotypes. N, sample number; ns, not statistically significant; ***p < 0.001. (F) Tail bleeding time of 8- to 10-week-old mice as in panel C was recorded up to 6 min. N, sample number; **p < 0.01.

### *Rbm8a* Knockout Impairs Megakaryocyte Differentiation

To explore how *Rbm8a* participates in platelet production, we first examined megakaryocytes in sections of bone marrow and spleen from 9-week-old mice. Hematoxylin and eosin (HE) staining revealed that *Rbm8a* knockout significantly reduced the average size of megakaryocytes in the bone marrow although the number of immature megakaryocytes increased (Figure 3A, HE, dot and bar graphs). IHC staining revealed that the level of von Willebrand factor (vWF), a marker for mature megakaryocytes, was notably reduced, suggesting that megakaryocyte differentiation was incomplete in *Rbm8a*KO^MK^ mice (Figure 3A, IHC). In the spleen, *Rbm8a*KO^MK^ megakaryocytes formed clumps (Figure 3B, HE, yellow dashed line), so that their number was greatly increased (bar graph) and, however, their size was reduced (dot graph). Accordingly, IHC staining revealed reduced vWF in splenic megakaryocytes, indicating impairment of their differentiation pathway (Figure 3B, IHC). Moreover, the relatively high nuclear-to-cytoplasmic ratio for *Rbm8a*KO^MK^ megakaryocytes reflected their immaturity. The observed accumulation of megakaryocytes in the spleen constituted evidence of splenomegaly in *Rbm8a*KO^MK^ mice.

**Figure 3.**
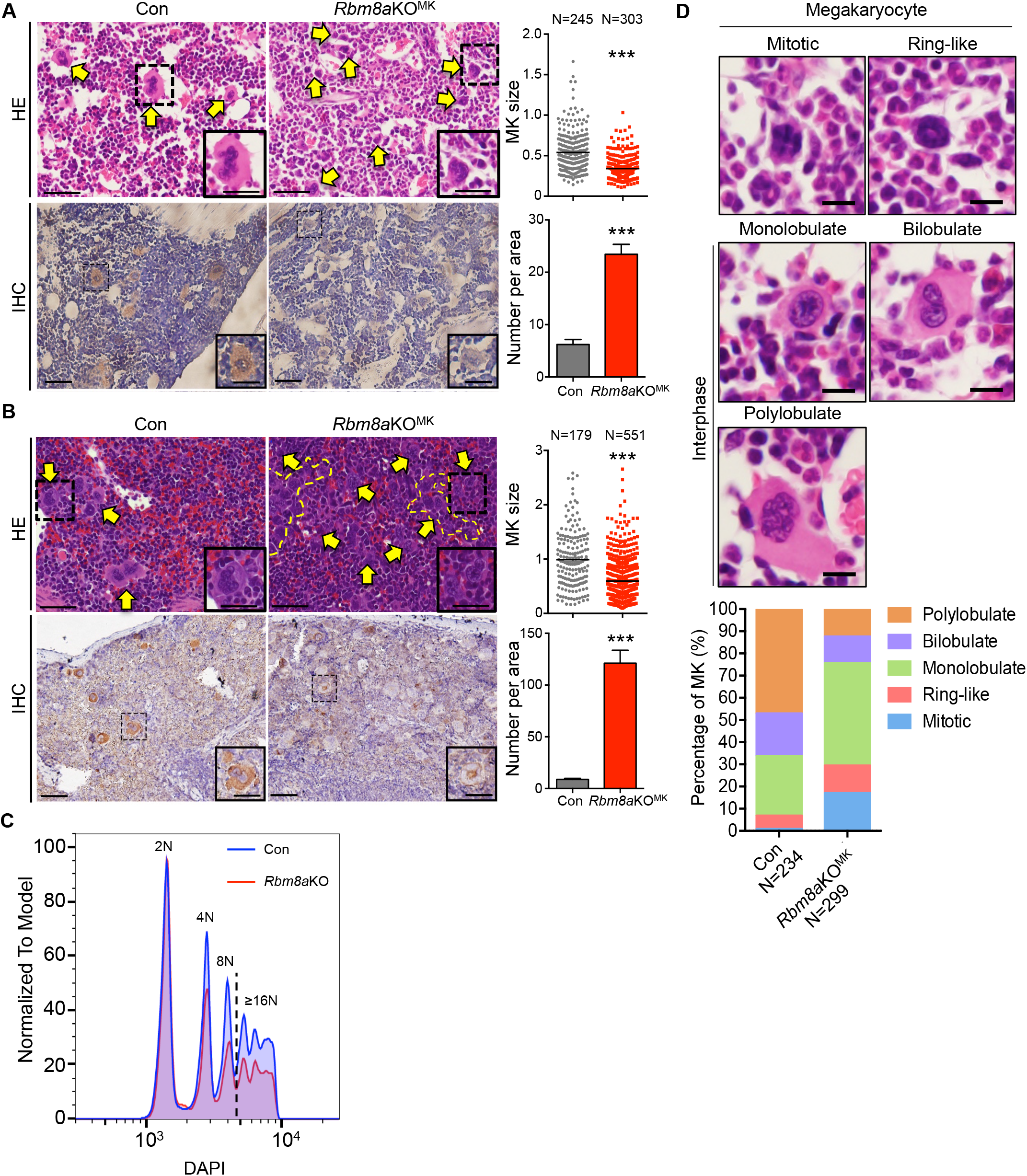
*Rbm8a* Knockout Impairs Megakaryopoiesis. Hematoxylin and eosin (HE) staining or immunohistochemical (IHC) staining (anti-vWF) of sections of bone marrow (A) and spleen (B) from 9-week-old Con and *Rbm8a*KO^MK^ mice. For both panels, yellow arrows indicate megakaryocytes. In panel B, yellow dashed lines indicate a cluster of megakaryocytes. A selected megakaryocyte in the dashed-line square is magnified in the inset. Scale bars represent 50 μm (insets, 20 μm). Dot plots show the size of megakaryocytes measured from HE staining (1.0= 500 μm^2^); for each, N indicates the number of cells from five mice of each genotype (mean ± SEM) that were measured. The bar graph shows the number of megakaryocytes in a 0.15 mm^2^ area; mean ± SEM was calculated from five areas each. *** p < 0.001. MK, megakaryocyte. (C) DNA histogram analysis of bone marrow of Con and *Rbm8a*KO^MK^ mice that were stained using DAPI. Each stacked-column bar shows the ploidy distribution of CD41^+^ cells. (D) Nuclear morphology of Con and *Rbm8a*KO^MK^ megakaryocytes was analyzed from panel A (HE). Each stacked-column bar shows the proportion (%) of each nuclear shape. Representative images of the different nuclear shapes were obtained from Con mice. Bar scale, 10 μm.

Accumulation of immature megakaryocytes indicated that *Rbm8a* deficiency impaired megakaryocyte maturation. Because polyploidization is a part of the megakaryocyte differentiation process, we measured DNA content of *Rbm8a*KO^MK^ megakaryocytes. Flow cytometry revealed a ∼1/3 reduction in the fraction of megakaryocytes with higher ploidy in *Rbm8a*KO^MK^ mice, indicating that *Rbm8a* knockout interfered with megakaryocytic polyploidization (Figure 3C). Along with an increase in ploidy, differentiated megakaryocytes display a single polylobulated nucleus (Mazzi et al., 2018). Therefore, we examined nuclear lobulation of megakaryocytes. HE staining revealed that ∼50% of control megakaryocytes were polylobulated and nearly 20% bilobulated, whereas ∼50% of *Rbm8a*KO^MK^ megakaryocytes were monolobulated (Figure 3D). Moreover, *Rbm8a* knockout likely prolonged mitosis and/or caused mitotic arrest (Figure 3D, blue). Together, these results indicated that *Rbm8a* is essential for megakaryocyte differentiation.

### Y14 Depletion Impairs Endomitosis of Megakaryocytes

The human erythroleukemic (HEL) cell line behaves as pluripotent hematopoietic precursors and can be induced to differentiate into megakaryocytes upon treatment with phorbol 12-myristate 13-acetate (PMA) (Papyyannopoulou et al., 1983). Therefore, we explored how Y14 participates in megakaryocyte differentiation of HEL cells. PMA induced the expression of CD41/GPIIb and CD61/GPIIIa, indicating that the cells had undergone differentiation to yield megakaryocytes (Figure 4A, cp. bars 1 and 3). Depletion of Y14 by lentiviral transduction of a Y14-targeting short hairpin RNA (shRNA) reduced the expression of CD41 and CD61 in PMA-treated HEL cells (Figure 4A, cp. bars 3 and 4), indicating that Y14 is essential for megakaryocyte differentiation. Next, we evaluated the role of Y14 in polyploidization. In particular, PMA increased the 8N population from 7% to 23% (Figure 4B). Depletion of Y14 in PMA-treated HEL cells substantially abolished the 4N/8N peaks and intriguingly induced a peak between 2N and 4N (Figure 4B, shY14_PMA, arrow); that peak was not observed without PMA treatment (shY14_mock). Moreover, reduced incorporation of ethynyl deoxyuridine (EdU) in Y14-depleted/PMA-treated cells argued that this peak was not the result of prolonged S phase (Figure S1). Immunoblotting and IHC revealed an increased level of phosphorylated histone 3 (phosphor-H3) in Y14-depleted HEL cells and bone-marrow megakaryocytes (Figure 4C, and Figure S2, upper). Therefore, Y14 depletion likely caused cell-cycle arrest as previously reported (Ishigaki et al., 2013; Lu et al., 2017), and impaired endomitosis as evident by reduced DNA synthesis. Because the features of *Rbm8a*KO^MK^ megakaryocytes, including reduced ploidy and nuclear lobe number and increased phospho-H3 and γH2AX (Figure S2, lower), were reminiscent of *Cdc20* null megakaryocytes (Trakala et al., 2015), we examined Cdc20 in Y14-depleted HEL cells. Immunoblotting revealed that Y14 knockdown reduced the level of Cdc20 by ∼60% in PMA-treated HEL cells (Figure 4C), further supporting a critical role for Y14 in endomitosis.

**Figure 4.**
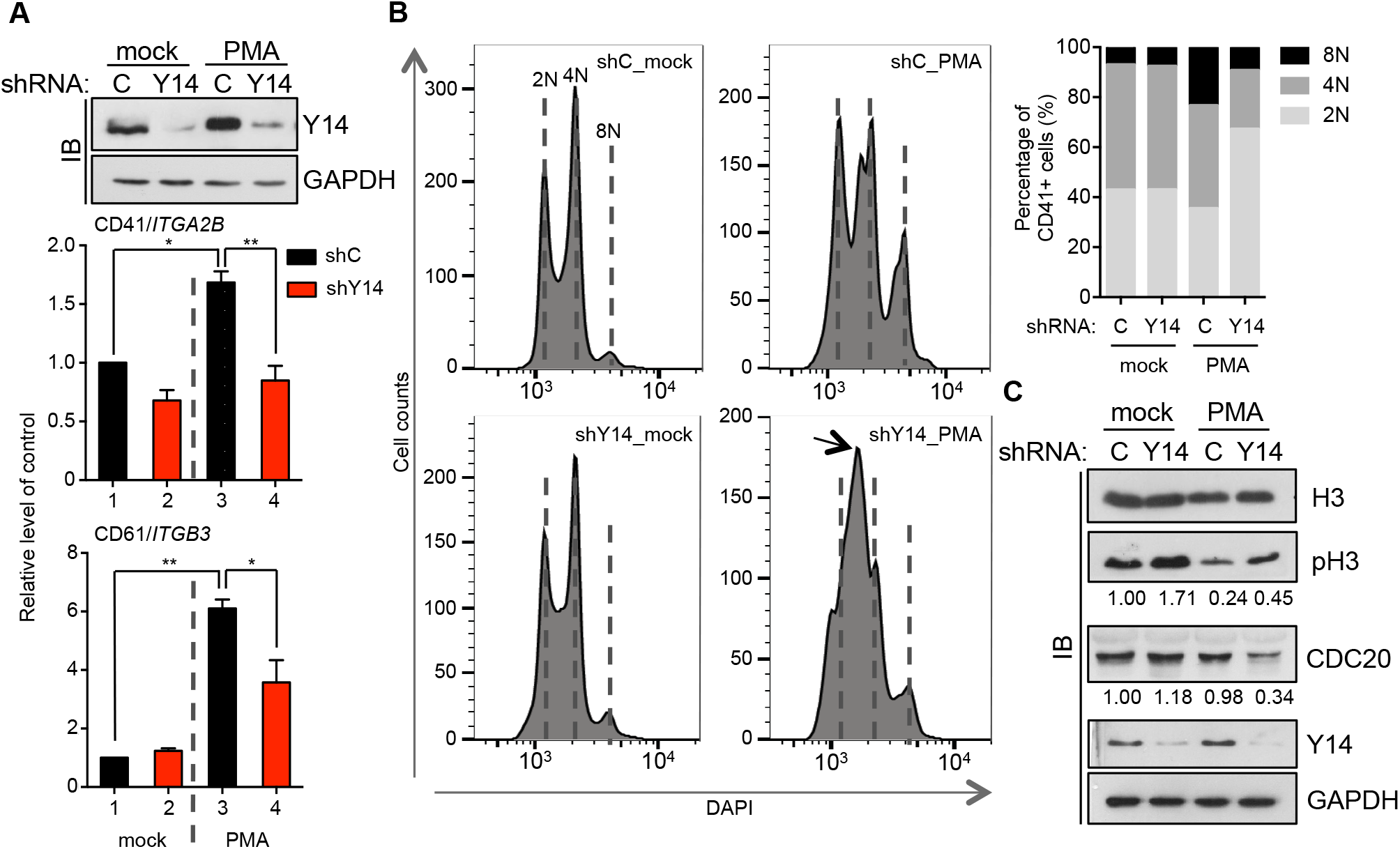
Depletion of Y14 Impairs Polyploidization via Cell Cycle Arrest. (A) HEL cells were transduced with lentivirus expressing shY14 (+) or control shRNA (–) for 48 h at a multiplicity of infection of 5, followed by mock-treated or PMA (5 nM) treatment for 3 days. RT-qPCR was performed to detect CD41/*ITGA2B* and CD61/*ITGB3*; experiments were performed in triplicate (**p < 0.01). Immunoblotting (IB) was performed using antibodies against Y14 and GAPDH. (B) HEL cells were transduced and treated as in panel A. DNA histogram analysis was performed as in Figure 3C. The stacked-column bar shows ploidy distribution; a peak between 2N and 4N (arrow) in the PMA-treated/Y14-depleted (shY14_PMA) cells was quantified as 2N. (C) HEL cells were transduced and treated as in panel A. Immunoblotting was performed using antibodies as indicated. The number below each lane of phospho-H3 (pH3) and CDC20 indicates their relative expression level (lane 1 was set to 1). Experiments were performed in triplicate.

### Y14 Deficiency Compromises Megakaryocyte Maturation and Disrupts Platelet Function

Next, we evaluated whether Y14 depletion in HEL cells alters gene expression profiles during megakaryocyte differentiation. We performed RNA-seq in control or Y14-depleted HEL cells without or with PMA treatment. Gene Ontology enrichment analysis of genes that were differentially expressed by >2 fold revealed that the top 10 groups of genes induced by PMA were functionally linked to granule formation, Ras and kinase signaling, cell adhesion, receptor signaling pathways, cytoskeletal reorganization, and endoplasmic reticulum stress response (Figure 5A, upper). Analysis of Y14-dependent functional groups also identified factors involved in the endoplasmic reticulum stress response and Fc receptor signaling (Figure 5A, lower); in each group, ∼70% of Y14-depletion downregulated genes were PMA-inducible (Figure 5B). Reverse transcription and quantitative PCR (RT-qPCR) revealed that PMA induced the expression of several endoplasmic reticulum stress response factors, such as CXCL8/IL-8, HSPA13, and CREB1, in HEL cells, and Y14 depletion compromised this induction (Figure 5C, left). Analogously, the expression of Fc receptor signaling factors, such as NFATC1, PRKCD, and FcGR2A, which are involved in platelet activation, was also induced by PMA and the induction was blunted upon Y14 depletion (Figure 5C, right). This result indicated that Y14 is also important for megakaryocyte maturation and even platelet function via its role in gene regulation. Therefore, we examined the expression of several platelet surface glycoproteins, which function to initiate platelet activation and aggregation, in *Rbm8a*KO^MK^ mice. Although *Rbm8a* knockout drastically reduced the platelet count, a residual amount of platelets was sufficient for flow cytometry. The result revealed a substantial reduction in the expression of the glycoproteins GP-VI and GP-IX and integrins α2bβ3 and CD41 in *Rbm8a*KO^MK^ platelets (Figure 5D). Moreover, we treated whole blood with the protease-activated receptor 4 (PAR4) agonist peptides, which induce platelet activation (Kahn et al., 1998). Flow cytometry revealed that *Rbm8a*KO^MK^ platelets exhibited a severe defect in inside-out activation of β3 integrin (JON/A) and α-granule secretion of fibrinogen, vWF, and P-selectin after treatment with a PAR4 agonist (Figure 5E). Thus, *Rbm8a* knockout not only impaired polyploidization but also disrupted surface glycoprotein expression and activation of platelets.

**Figure 5.**
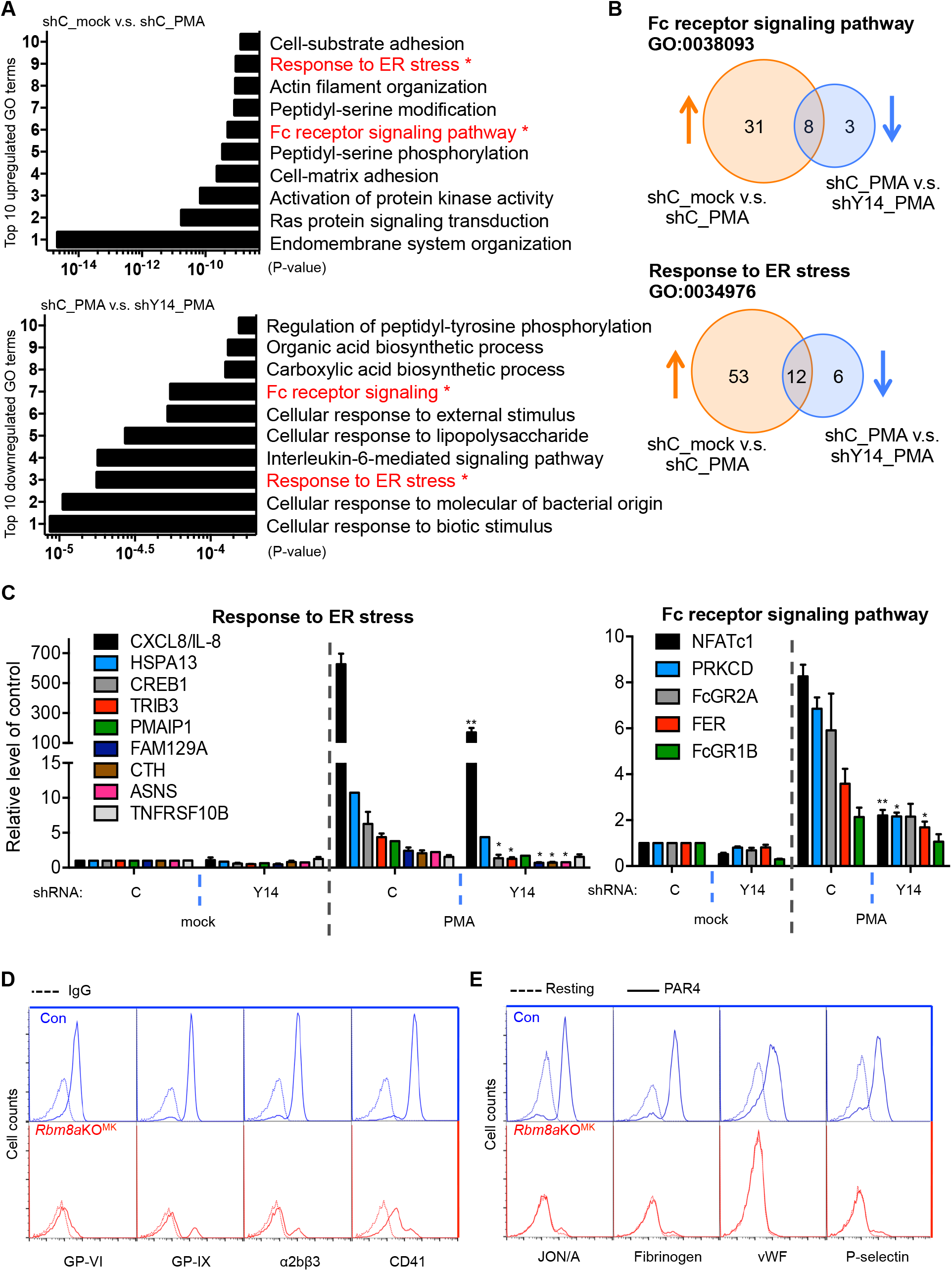
*Rbm8a* Deficiency Impairs Megakaryocyte-Specific Gene Expression and Platelet Function. (A) RNA-seq analysis of HEL cells that were transduced and PMA-treated as in Figure 4A. Gene Ontology (GO) enrichment analysis was performed on differentially expressed genes that exhibited a fold-change of >2 and a *p*-value of <0.05. Bar graphs show the top 10 largest changes in each comparison group (shC_mock vs. shC_PMA, and shC_PMA vs. shY14_PMA). Red highlights represent the categories changed upon PMA induction and Y14 depletion. ER, endoplasmic reticulum. (B) Venn diagrams showing the overlaps between PMA-induced genes (orange) and genes for which expression was reduced upon Y14 depletion in differentiated HEL cells (PMA-treated; blue) in the two indicated categories of genes. (C) RT-qPCR analysis of the indicated genes in HEL cells that were transfected with shRNA and then mock-treated or treated with PMA as in panel A. The experiments were performed at least three times. *p < 0.05, **p < 0.01. (D) Whole blood of Con and *Rbm8a*KO^MK^ mice was collected and stained for GP-VI, GPIX, α2bβ3, CD41 or IgG (dashed lines). Platelets were gated by forward scattering/side scattering characteristics. Representative flow cytometry profiles showing the indicated glycoproteins or integrins. (E) Blood was mock-treated (resting; dashed lines) or treated with PAR4 (solid lines) and subsequently stained with an antibody against JON/A (phycoerythrin), fibrinogen, vWF, or P-selectin. Flow cytometry profiles of platelet glycoproteins are shown as in panel D.

### Inhibition of p53 Reverses Differentiation Defects of *Rbm8a*KO^MK^ Megakaryocytes

Because Y14 deficiency increases the level of p53 protein (Lu et al., 2017; Mao et al., 2016), we suspected that *Rbm8a* knockout would increase the level of p53 in megakaryocytes, which would hence interfere with megakaryocyte differentiation. IHC staining of bone-marrow sections using anti-p53 revealed a higher level of p53 in *Rbm8a*KO^MK^ megakaryocytes than in control megakaryocytes (Figure 6A). To explore how the possible Y14-p53 regulatory axis influences megakaryocyte differentiation, we first examined the expression of Y14 and p53 in HEL cells following PMA treatment. Immunoblotting revealed a decrease in p53 and an increase in Y14, indicating inverse expression of Y14 and p53 during megakaryocyte differentiation (Figure 6B). Knockdown of Y14 increased the level of p53 in PMA-treated HEL cells (Figure 6C), consistent with that observed in bone-marrow megakaryocytes (Figure 6A). Because p53 negatively regulates *Cdc20* expression (Banerjee et al., 2009), we postulated that increased p53 would cause a reduction in Cdc20 in Y14-depleted HEL cells. Therefore, we treated Y14-depleted HEL cells with the p53 inhibitor pifithrin-α (PFT-α). Immunoblotting revealed that PFT-α restored Cdc20 level and reduced phospho-H3 accumulation in a dose-dependent manner (Figure 6D). PFT-α treatment shifted the promiscuous peak between 2N and 4N to 2N but failed to increase the ploidy (Figure 6E), suggesting that p53 inactivation at least partially restored the cell cycle. Next, we evaluated whether p53 inhibition could promote *ex vivo* differentiation of Y14-deficient megakaryocytes. *Rbm8a*KO^MK^ megakaryocytes in the fetal liver section exhibited a higher level of p53 and γH2AX (Figure S3), as observed in bone marrow (Figures 6A and Figure S2). *Rbm8a*KO^MK^ megakaryocyte progenitors derived from fetal liver stroma were smaller than those of the control (Figure 6F, mock treatment). PFT-α treatment restored their average size and CD41 level (Figure 6F, bar graphs).

**Figure 6.**
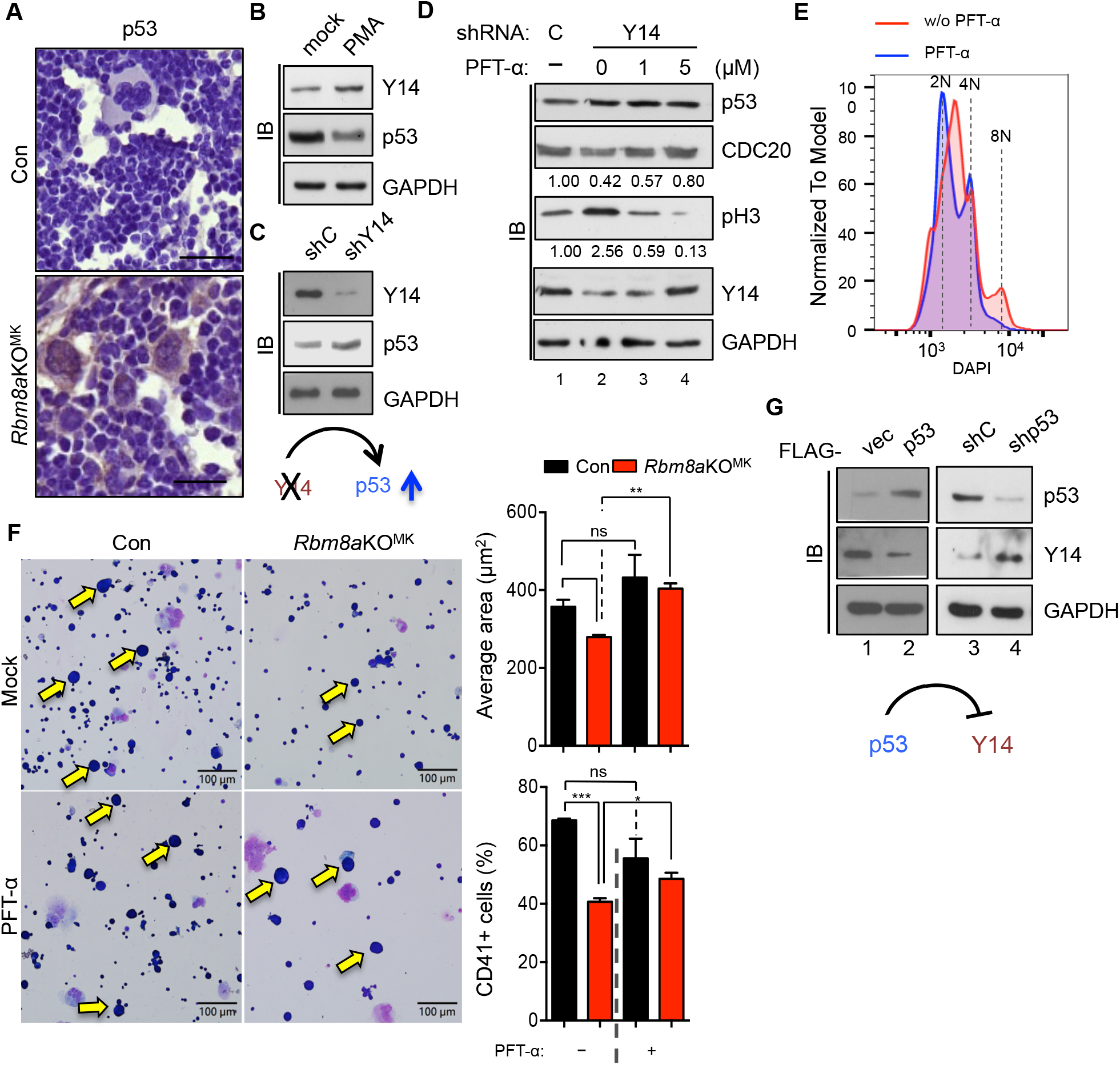
p53 Inhibition Reverses the Differentiation Defects of *Rbm8a*KO^MK^Megakaryocytes. (A) Bone-marrow sections as in Figure 1D from five mice of each group were subjected to immunohistochemical staining using anti-p53; bar scale, 20 µm. (B) HEL cells were mock-treated or PMA-treated for 3 days. Panels B–D, G: Immunoblotting (IB) was performed with the indicated antibodies. Experiments were performed in triplicate. (C) HEL cells were transduced with control or shY14-lentivirus. (D) HEL cells were transduced with control or shY14-lentivirus followed by mock (–) or PFT-α (1 or 5 µM) treatment. The relative levels of CDC20 and phospho-H3 (pH3) are indicated below the respective blots; lane 1 was set to 1. (E) HEL cells were transduced with shY14-lentivirus and treated with PMA followed by mock or PFT-α treatment. DNA histogram analysis was performed as in Figure 3C. (F) Primary liver cells from Con or *Rbm8a*KO^MK^ embryos on embryonic day 14.5 containing megakaryocyte precursors were precultured in the presence of thrombopoietin. Enriched megakaryocytes were mock-treated or treated with PFT-α. Yellow arrows indicate megakaryocytes; bar scale, 100 µm. Bar graphs show the average size (µm^2^) of >350 cells (upper) and the percentage of CD41^+^ (lower) in each group. The mean ± SEM was measured; *p < 0.05, **p < 0.01, ***p < 0.001; ns, not significant. (G) HEL cells were transfected with the empty (vec) or p53-expressing vector (lanes 1–2), or shC or shP53-expressing lentivirus (lanes 3–4). The cartoon shows that p53 may suppress Y14 expression during megakaryocyte differentiation.

The above result unexpectedly revealed that a high dose of PFT-α could increase the cellular level of Y14 (Figure 6D, lane 4). Therefore, we transiently overexpressed or knocked down p53 in HEL cells to evaluate Y14 expression. Overexpression of p53 suppressed the expression of Y14, possibly at the transcriptional level (Figure 6G, and Figure S4). Therefore, p53 may restrict Y14 expression in megakaryoblasts. Upon induction of differentiation, the consequent downregulation of p53 allows an increase of Y14, which is essential for polyploidization and maturation of megakaryocytes.

### p53 Knockout Rescues Platelet Production in *Rbm8a*KO^MK^ Mice

In light of the *in vitro* observation that p53 inhibition could in part reverse the differentiation defects of *Rbm8a*KO^MK^ megakaryocytes, we had attempted to evaluate the effect of PFT-α in mice. Following daily intravenous injection of PFT-α for several days, mice unfortunately died of excessive bleeding. Therefore, we turned to genetically ablate p53 in *Rbm8a*KO^MK^ mice. Pf4^Cre/+^;*Rbm8a*^f/+^ mice were crossed with *Trp53*^f/f^. The resulting *Rbm8a*^f/+^;*Trp53*^f/+^ mice were then crossed with *Trp53*^f/f^ to generate *Rbm8a*^f/+^;*Trp53*^f/f^. Subsequently, the use of Pf4^Cre/+^;*Rbm8a*^f/+^;*Trp53*^f/+^ and *Rbm8a*^f/+^;*Trp53*^f/f^ mice generated all genotypes needed for the analysis (Figure 7A and Figure S5A for concise and complete diagrams, respectively); the *Trp53* genotypes of the offspring were confirmed by PCR analysis (Figure S5B). Moreover, using PCR and Sanger sequencing, we confirmed Cre-mediated excision of floxed *Trp53* sequences from bone marrow cells of floxed allele-bearing mice (Figure S5C, D).

**Figure 7.**
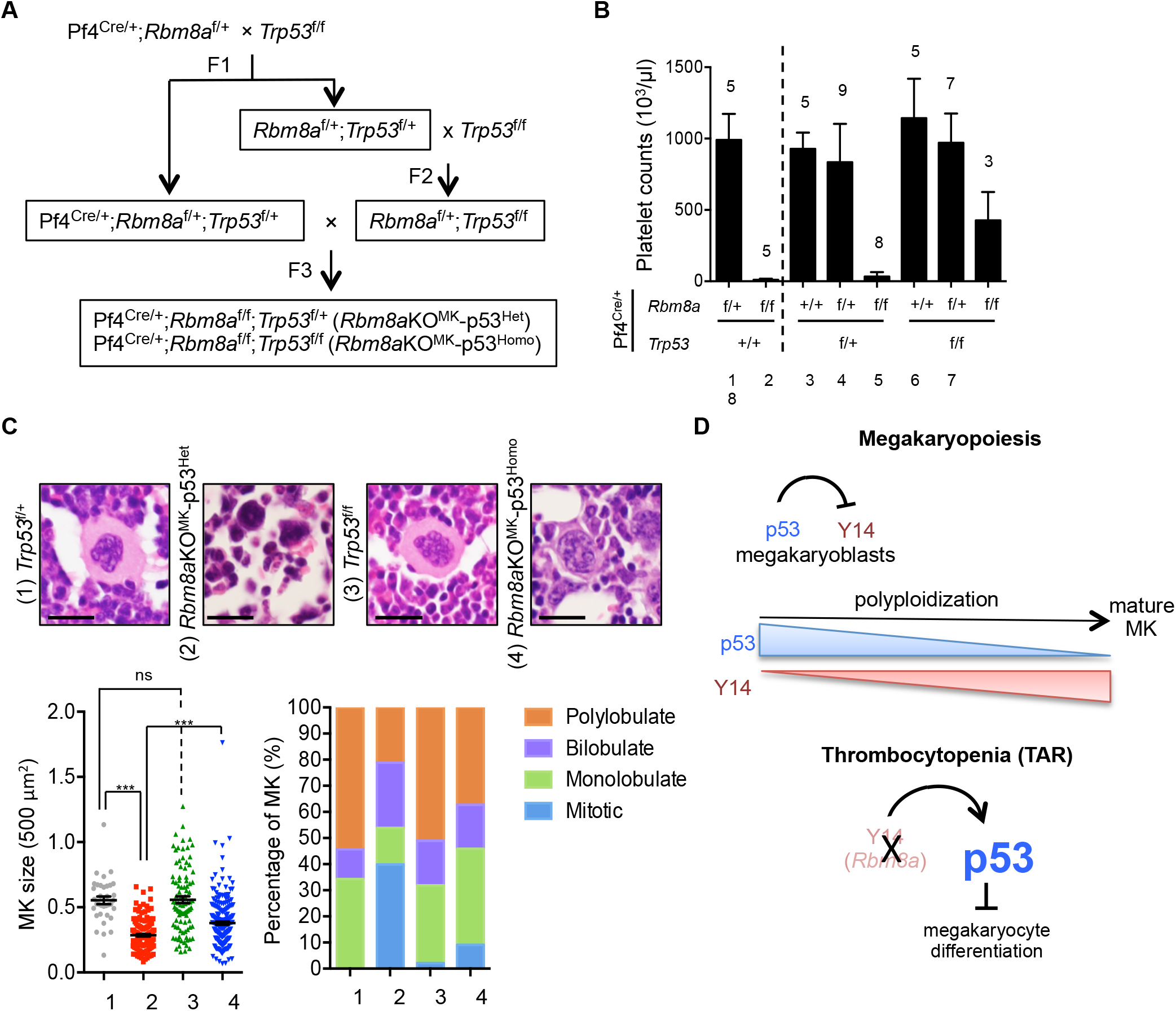
*Trp53* Knockout Partially Restores Platelet Production in *Rbm8a*KO^MK^ Mice. (A) Schematic diagram showing the generation of megakaryocyte-specific *Rbm8a*/*Trp53* double-knockout mice. A more detailed diagram is shown in Figure S5A. (B) Hematological analysis of blood from 6- to 8-week-old mice of the indicated genotypes. Bar graph shows platelet counts as in Figure 2E; the number of mice was indicated above each bar. Since most of *Rbm8a*KO^MK^-p53^Homo^ mice died at 3-5 weeks of age, data were acquired for only three mice. (C) HE staining of bone marrow sections from *Trp53*^f/+^, *Trp53*^f/f^, *Rbm8a*KO^MK^-p53^Het^ and *Rbm8a*KO^MK^-p53^Homo^ mice (see Figure S7); magnified images of representative cells are shown (bar scale, 20 µm). Stacked-column bar graph shows the proportion (%) of cells with different morphological features. Dot plots show the size of megakaryocytes measured from HE staining; n.s. not significant; ***p < 0.001. Stacked-column bar graph shows the proportion (%) of cells with different morphological features. For each genotype, 100∼200 megakaryocytes were measured. (D) Model showing the Y14/p53 regulatory circuit in megakaryocyte differentiation and its dysregulation in TAR syndrome. p53 may suppress Y14 expression in megakaryoblasts. Upon induction of differentiation, p53 is downregulated, whereas Y14 is upregulated and participates in polyploidization and megakaryocyte maturation. Thrombocytopenia (TAR): *Rbm8a* deficiency or Y14 depletion activates p53, causes cell-cycle arrest, and impairs megakaryocytic gene expression, leading to a blockade of megakaryopoiesis. Treatment with p53 inhibitors or *Trp53* knockout restores megakaryocyte differentiation and platelet production, respectively.

Pf4^Cre/+^;*Rbm8a*^*f/f*^;*Trp53*^*+/+*^ (namely *Rbm8a*KO^MK^-p53^WT^) had very low platelet counts, as expected (Figure 7B, lane 2). Knockout of one *Trp53* allele in *Rbm8a*^+/+^ or *Rbm8a*^f/+^ mice did not affect platelet production (Figure 7B, lanes 3 and 4), whereas knockout of two alleles minimally increased the platelet count (lanes 6 and 7), which may have resulted from slightly enhanced megakaryocyte differentiation due to p53 deficiency (Apostolidis et al., 2012). Pf4^Cre/+^;*Rbm8a*^f/f^;*Trp53*^f/f^ (*Rbm8a*KO^MK^-p53^Homo^), however, acquired tumors in lymph nodes and most of them died at 3-5 weeks old regardless of gender (Figure S6 and Table S3). Nevertheless, blood analysis of the three surviving mice showed a partial restoration of the platelet count (Figure 7B, lane 8), supporting our hypothesis that p53 inactivation/deficiency could in part rescue the defects caused by *Rbm8a* knockout. These three mice died after blood collection and we found that their lymph nodes also bore tumors. Knockout of one *Trp53* allele was unable to restore platelet production of *Rbm8a*KO^MK^ (Figure 7B, lane 5, Pf4^Cre/+^;*Rbm8a*^f/f^;*Trp53*^f/+^; *Rbm8a*KO^MK^-p53^Het^). These mice were in general alive > 2 months, although two out of eight still developed tumors in lymph nodes. Due to the early death of *Rbm8a*KO^MK^-p53^Homo^, we sacrificed 3-4 *Rbm8a*KO^MK^-p53^Het^ and *Rbm8a*KO^MK^-p53^Homo^ mice to examine their bone marrow using HE staining. The control *Trp53*^f/+^ or *Trp53*^f/f^ megakaryocytes showed normal morphological features (Figure S7 for a representative field of cells; Figure 7C for a magnified image). The majority of *Rbm8a*KO^MK^-p53^Het^ megakaryocytes had a hypolobated or condensed nucleus, indicating cell cycle arrest and impaired polyploidization (Figure 7C). Homozygous knockout of p53 substantially reduced the population of cells that exhibited mitotic arrest, and hence *Rbm8a*KO^MK^-p53^Homo^ megakaryocytes exhibited monolobular and even polylobular nuclei (Figure 7C). However, their small cytoplasmic volume and accumulation in the bone marrow indicated immaturity (Figure 7C).

Together, our results demonstrated the role of *Rbm8a/*Y14 in megakaryocyte differentiation and platelet production, which involves p53. Knockout of p53 in *Rbm8a*-deficient megakaryocytes could partially restore megakaryopoiesis and platelet counts. Moreover, we uncovered the potential Y14-p53 regulatory circuit in the process of megakaryopoiesis.

## DISCUSSION

In this study, we show that *Rbm8a* deficiency resulted in a blockade of megakaryocyte differentiation partially through cell cycle disruption (*i.e*., phospho-H3 accumulation) and p53 activation (Figures 6A, S2 and S3). A similar result was observed in Y14-depleted HEL cells (Figure 6D). We further found that Y14 depletion reduced the level of mitotic regulators Cdc20 (Figure 4C) as well as CDK1 and cyclin B1; the latter two were revealed by transcriptomic analysis and then confirmed by immunoblotting (Figure S8). These results supported the notion that Y14 deficiency causes cell cycle dysregulation. It should be noted that HEL cells express a high level of p53 in which there is a mutation (M133K) in the DNA-binding domain (Zhao et al., 2012). Nevertheless, this mutant p53 retained partial transactivation activity (Figure S9), and its expression level was further increased upon Y14 depletion (Figure 6C and 6D). p53 activation may thus contribute to cell cycle arrest and decreased ploidy in Y14-depleted megakaryocytes. PFT-α treatment restored Cdc20 expression and reduced the level of phospho-H3 (Figure 6D), and shifted the intriguing peak between 2N and 4N to 2N (Figure 6E). Therefore, p53 inhibition may have reactivated cell cycle progression of Y14-depleted HEL cells but was insufficient to promote endomitosis. Perhaps abundant mutant p53 that was not fully inactivated by PFT-α interfered with endomitosis. Additionally, the observed induction of cell cycle arrest via Y14 deficiency may be independent of p53, at least in part. Nevertheless, PFT-α treatment could restore megakaryocyte differentiation *ex vivo*, as evidenced by the increased cell size and CD41 expression (Figure 6F).

Previous studies have revealed that haploinsufficiency of the exon junction complex core components including *Rbm8a* causes microcephaly, and complete knockout of p53 could rescue neurodevelopmental defects (Mao et al., 2015; Mao et al., 2016; Silver et al., 2010). Here we show that heterozygous *Rbm8a* knockout had no effect on platelet counts, whereas complete loss of *Rbm8a* abolished platelet production and function. Our result echoes the scenario of TAR syndrome, in which one *RBM8A* allele is deleted in concert with mutations (and consequent downregulation) of another allele (Albers et al., 2013), indicating that drastic reduction of Y14 impairs megakaryocyte differentiation. Although both genders exhibited internal hemorrhage and splenomegaly, weaning-age male *Rbm8a*KO^MK^ had more severe symptoms such as intestinal bleeding and, ultimately, death. It has been reported that Pf4-Cre-mediated recombination minimally occurs in circulating CD45^+^ cells and the distal intestine (Pertuy et al., 2015). Perhaps *Rbm8a* knockout is harmful to rapidly proliferating intestinal epithelial cells, but why such an effect was prominent in males is unclear. Our results revealed that knockout of one p53 allele in *Rbm8a*KO^MK^ was not sufficient to restore the platelet count. Complete loss of p53 may resume cell cycle progression in *Rbm8a* deficient megakaryocytes as evidenced by their nuclear lobulation (Figure 7). Perhaps p53 reduction was unable to completely restore additional defects, such as megakaryocyte-specific gene expression, caused by *Rbm8a* knockout. Therefore, *Rbm8a*KO^MK^-p53^Homo^ megakaryocytes were not fully mature, which compromised the platelet production. Ablation of p53 yet induced tumors in lymph nodes of *Rbm8a* knockout mice, leading to early death (Figure S6 and Table S3). A high frequency of lymphomas has been observed in *Trp53*-null mice (Donehower et al., 1995). Because Y14 is also essential for maintaining genome stability (Chuang et al., 2019), concomitant loss of *Rbm8a* and p53 may likely increase the incidence of cancer.

Our results reveal that p53 can suppress Y14 expression (Figure 6G). Upon PMA induction, Y14 and p53 were respectively upregulated and downregulated in HEL cells (Figure 6B). Therefore, it is probable that p53 restricts Y14 expression in undifferentiated megakaryocytes. It should be noted that although Y14 depletion increases p53 expression, overexpression of Y14 or any other exon-junction components promotes p53 mRNA translation (Lu et al., 2017). Therefore, we excluded the possibility that the increase of Y14 results in p53 downregulation during megakaryocyte differentiation and assumed that factors other than Y14 suppress p53 expression. Nevertheless, p53 is activated under Y14-deficient conditions, such as TAR syndrome, leading to a blockade of megakaryopoiesis. Therefore, the regulatory network of Y14 and p53 likely modulates megakaryopoiesis, and its dysregulation results in platelet disorders (Figure 7D).

*Rbm8a* deficiency results in a blockade of megakaryopoiesis via p53 activation. p53 activation has been observed in several congenital syndromes, such as Diamond-Blackfan anemia and 5q^−^ syndrome, which exhibit ribosomal mutations or defects in ribosomal biogenesis (Deisenroth et al., 2016). These diseases are characterized by macrocytic anemia and/or hypolobulated micromegakaryocytes, indicating defective erythropoiesis or megakaryopoiesis. Knockdown of *RPL11* (associated with Diamond-Blackfan anemia) in human CD34^+^ hematopoietic stem/progenitor cells inhibits cell proliferation and erythropoiesis, and such defects can be reversed by p53 knockdown (Moniz et al., 2012), which is similar to the scenario for the Y14-p53 circuit in megakaryopoiesis. In addition to ribosomal factors, certain cell cycle or DNA damage repair factors are also involved in megakaryopoiesis. As described above, the phenotype of *Rbm8a* knockout mice was similar to that of *Cdc20* knockout, including a blockade of polyploidization, nuclear hypolobulation, and reduced platelet production (Trakala et al., 2015). Y14 depletion reduced the level of Cdc20 in HEL cells (Figure 4C). Knockdown/inhibition of p53 restored the level of Cdc20 and consistently resumed the cell cycle in HEL cells (Figures 6D and 6E), suggesting that the Y14-p53 circuit acts upstream of Cdc20 during megakaryopoiesis. Finally, it is interesting to note that thrombopoietin can trigger the non-homologous end-joining pathway in hematopoietic stem cells perhaps to ensure chromosomal integrity (de Laval et al., 2013). Moreover, knockout of Fanconi anemia-associated *Fanca* results in defective megakaryopoiesis (Pawlikowska et al., 2014), further suggesting that the DNA damage response network is important for preventing DNA damage accumulation through endomitotic cycles. Thus, whether the role of Y14 in DNA damage repair and genome stability maintenance (Chuang et al., 2019) also contributes to megakaryocyte differentiation remains to be deciphered.

Overall, our results show that an increase in p53 accounts for impaired megakaryocyte differentiation in *Rbm8a*KO^MK^ mice, which can be rescued by inhibition or knockout of p53. Therefore, TAR syndrome represents a type of p53 activation that can cause developmental defects, and the use of p53 inhibitors may possibly be a therapeutic approach in the future.

## EXPERIMENTAL PROCEDURES

### Generation of Megakaryocyte-Specific Knockout Mice

Conditional allele of *Rbm8a* (*Rbm8a*^f/f^) was created by insertion of *loxP* sites flanking its exon 2 and 6 using the CRISPR/Cas9 system (Chuang et al., 2019). *Rbm8a*^f/f^ mice were then crossed with Pf4-Cre^+/-^ mice (Tiedt et al., 2007). We first generated heterozygous mice for Cre and floxed *Rbm8a* (Pf4-Cre^+/-^;*Rbm8a*^f/+^, Con) that were then bred to *Rbm8a*^f/f^ to produce *Rbm8a* megakaryocyte/platelet-specific knockout (Pf4-Cre^+/-^;*Rbm8a*^f/f^ *Rbm8a*KO^MK^) mice (Figure 1C). These mice were born at the expected Mendelian frequency (Table S1). Genotyping was conducted by PCR of toe lysate (primers are listed in Table S4). To generate megakaryocyte/platelet-specific *Rbm8a* and *Trp53* (p53) knockout, we crossed *Rbm8a*KO^MK^ mice with *Trp53*^F2-10/F2-10^ (Jonkers et al., 2001) (hereafter termed as *Trp53*^f/f^). In brief, Pf4-Cre^+/-^;*Rbm8a*^f/+^ were mated with *Trp53*^f/f^ mice to generate Pf4-Cre^+/-^;*Rbm8a*^f/+^;*Trp53*^f/+^. Meanwhile, *Rbm8a*^f/+^;*Trp53*^f/+^ mice were bred with the *Trp53*^f/f^ mice to generate *Rbm8a*^f/+^;*Trp53*^f/f^. Next, Pf4-Cre^+/-^;*Rbm8a*^f/+^;*Trp53*^f/+^ were bred with *Rbm8a*^f/+^;*Trp53*^f/f^ to generate Pf4-Cre^+/-^ mice with *Rbm8a* (^+/+^, ^f/+^ or ^f/f^) and *Trp53* (^f/+^ or ^f/f^) (Figure 7A). All experimental procedures involving animals were approved by the Institutional Animal Care and Use Committee, Academia Sinica (protocols 12-12-449 and 17-08-1104) and complied with the Ministry of Science and Technology, Taiwan. In the text, Pf4-Cre^+/-^ is abbreviated as Pf4-Cre. Note that male Pf4-Cre;*Rbm8a*^f/f^ died before sexual maturity, and male Pf4-Cre; *Trp53*^f/f^ had a high incidence of scrotal swelling possibly due to testicular teratoma, which restricted multiple mating.

### Histochemistry, Immunohistochemistry, and Immunofluorescence

Paraffin-embedded mouse spleen and femur (bone marrow) sections were prepared according to standard procedures. Deparaffinized and dehydrated sections were stained with HE for histology. Primary cells were smeared and dried on slides, and then fixed in methanol for 5 min. The samples were stained with Giemsa stain (1:20 dilution) for 15 min, and air-dried after rinse. For IHC, retrieved sections were sequentially incubated with monoclonal anti-Y14 (Abcam) or γH2AX (Millipore) polyclonal antibodies against vWF, p53 (Proteintech) or phospho-histone-H3 (pH3) (Cell Signaling Technology) overnight at 4°C and horseradish peroxidase–conjugated secondary antibody for 1 h at room temperature, followed by staining with DAB Quanto (Thermo Fisher). The nuclei were counter-stained with hematoxylin. The size and number of megakaryocytes were analyzed and quantified by using ImageJ software.

### Flow Cytometry

Blood samples were collected in heparinized tubes (365965, BD Biosciences) using submandibular bleeding into. Unstimulated or activated platelets were stained with fluorophore (FITC or PE)-conjugated antibodies against cell surface antigens for 15 min at room temperature, including monoclonal antibodies against GP-VI, GP-IX, α2bβ3, JON/A or P-selectin, and polyclonal antibodies against Fibrinogen or vWF (all above from EMFRET Analytics) or CD41 (BD Biosciences). Samples were analyzed by using an Attune NxT Flow Cytometer (Thermo Fisher).

### Cell Culture and Transfection, Megakaryocytic Differentiation, and p53 Inhibitor Treatment

HEL cells were grown in RPMI medium supplemented with penicillin-streptomycin and 10% fetal bovine serum (Gibco). For Y14 knockdown, HEL cells were transduced for 48 h with a Y14-targeting shRNA-expressing lentivirus at a multiplicity of infection of 5; shLuc served as the control. Megakaryocytic differentiation of HEL cells was archived by treatment with 5 nM PMA (Sigma) for 3 days. For p53 inhibition, HEL cells were treated with 5 μM PFT-α (Millipore) for 4 h prior to PMA induction; DMSO was used as control. Primary cells isolated from embryonic day 14.5 mouse fetal liver were grown in Dulbecco’s modified Eagle’s medium (Gibco) supplemented as above and with 100 ng/ml murine thrombopoietin (GenScript) for 3 days. Megakaryocytes were enriched by 1.5% and 3% BSA density-gradient centrifugation, and then cultured in the presence of thrombopoietin for another 3 days; for p53 inhibition, 5 μM PFT-α was added. To express or knock down p53, HEL cells were transfected with the p53 expressing vector (Lu et al., 2017) or transduced with p53-targeting shRNA lentivirus for 2 days.

### RNA Sequencing and Bioinformatics Analysis

Control shRNA or Y14-depleted HEL cells were mock-or PMA-treated for 3 days as described above. Two replicates of each treatment were subjected to RNA-seq. RNA was extracted by chloroform and selected by oligo-dT following the manufactory instructions. The libraries were constructed by KAPA mRNA HyperPrep Kit (KAPA Biosystems, Roche, Basel, Switzerland) and sequenced using Illumina Novaseq 6000 platform. A total of 53-81 million 150 bp paired-end reads were generated for each sample. Clean reads were generated by using Trimmomatic v.0.38 to filter low quality reads and remove adaptor sequences. Reads were aligned to GRCh38 database using HISAT2 (Kim et al., 2015; Sahraeian et al., 2017). Differential gene expression was analyzed using featureCounts (Liao et al., 2014). Differentially expressed coding genes were obtained using DESeq2 and defined as a fold-change of > 2 and a *p*-value of < 0.05 (Love et al., 2014). Gene Ontology Enrichment analysis was performed using clusterProfiler (3.10.1), showing top 10 differential expression categories of megakaryocytic differentiation-related (PMA-treated compared with mock-treated shLuc-HEL cells) or Y14-depletion-affected (shLuc-compared with Y14-depleted HEL cells after PMA treatment). A global expression profile of pre-defined curated gene sets and Gene Ontology gene sets from MSigDB were compared between control and Y14-depleted cells.

### Statistical Analysis

Statistical analyses were essentially performed using MedCalc statistic software (Ostend, Belgium) or Prism (GraphPad). Additional statistical analysis methods included two-tailed, unpaired and paired Student’s t test in this study. Data were presented as mean ± SEM. *P* values ≤0.05 were considered statistically significant.

### Giemsa Staining, Hematological Analysis, DNA Histogram Analysis, Tail Bleeding Assay, Plasmids, Immunoblotting, RT-PCR and RT-qPCR

The detailed procedures are in Supplemental Information.

## SUPPLEMENTAL INFORMATION

Supplemental Information includes Supplemental Experimental Procedures and nine figures.

## AUTHOR CONTRIBUTIONS

CHS and WJL performed experiments (CHS, Figures 1B, 3C, 3D, 4, 5A-C, 6, 7, and all supplemental figures; WJL, Figures 1D, 2, 3A, 3B, 5D, 5E, 7B), interpreted the data and contributed to manuscript preparation. WCK raised and maintained mice and performed the experiment of Figure 7B. RBY supervised the experiments. WYT designed the study, supervised the experiments, and wrote the manuscript.

## ACKNOWLEDGEMENTS

HEL cells and *Trp53*^f/f^ mice were respectively obtained from Tien-Shun Yeh and Li-Ru You (National Yang-Ming University, Taipei, Taiwan). We thank Kuan-Yang Hung and Chang-Yi Lin for generation of *Pf4*-Cre^+/-^;*Rbm8a*^f/+^ mice and initial characterization of HEL cells, respectively. We thank the Taiwan Animal Consortium funded by the Ministry of Science and Technology of Taiwan for technical support in hematological analysis. This study was supported by Academia Sinica Investigator Award (AS-IA-107-L04) to W.-Y.T.

