## Supplemental text for "The Y14-p53 Regulatory Circuit in Megakaryocyte Differentiation and Thrombocytopenia"

**SUPPLEMENTARY EXPERIMENTAL PROCEDURES**

**Giemsa Staining**

Primary cells from fetal liver stroma were smeared and dried on slides, and then fixed in methanol for 5 min. The samples were stained with Giemsa stain (1:20 dilution) for 15 min, and air-dried after rinse.

**Hematological Analysis**

Fresh blood was collected in K_2_EDTA tubes (BD Biosciences) and analyzed using ProCyte Dx® Hematology Analyzer (IDEXX Laboratories) according to manufacturer’s instruction.

**DNA Histogram Analysis**

Femoral and tibial bone marrow was isolated as described (Schulze, 2016). Bone-marrow cells were stained with FITC-conjugated rat anti-mouse CD41 or rat-IgG isotype control (both from BD Biosciences). Cells were washed in phosphate buffered saline (PBS) followed by fixation with 4% paraformaldehyde at room temperature for 1 h. After washing twice with PBS, cells were incubated with 5 μg/ml of DAPI and analyzed by flow cytometry (Trakala et al., 2015). HEL cells were stained with FITC-conjugated mouse anti-human CD41 (BD Biosciences), followed by flow cytometry analysis as above.

**Tail Bleeding Assay**

Mice were intraperitoneally anesthetized with a single dose of combined ketamine (100 mg/kg) and xylazine (15 mg/kg). Tail bleeding times were determined as described (Hoover-Plow et al., 2006). In brief, a distal 4-mm segment of the tail was amputated using a scalpel and immediately immersed in physiological saline maintained at 37˚C. The time for complete cessation of bleeding was recorded up to 6 min. The experiment was subsequently terminated in any mice that failed to stop bleeding.

**Plasmids and Cell Transfection**

HEK293 cells were grown in Dulbecco’s modified Eagle’s medium supplemented with penicillin-streptomycin and 10% fetal bovine serum. Gene specific shRNA-expressing lentiviral vectors were obtained from the RNAi Core Facility, Academia Sinica, including Y14 (5’-CGAGAGCATTCACAAACTGAA), p53 (5’-CACCATCCACTACAACTACAT), and control (C, 5’-GCGGTTGCCAAGAGGTTCCAT, which targeted luciferase). The pcDNA-FLAG-p53-expression vector was previously described (Lu et al., 2017). The p53 M133K mutant expression vector was generated by using PCR-based mutagenesis in the parent pCDNA-FLAG-p53 vector (Lu et al., 2017). For overexpression of FLAG-tagged p53 or M133K mutant, HEL cells were transfected with 4 μg of each FLAG-tagged vector by lipofectamine 2000 (Thermo Fisher Scientific) for 2 days.

**Immunoblotting**

Immunoblotting was performed as described (Lu et al., 2017), using monoclonal antibodies against GAPDH (Proteintech) and histone 3 (Cell Signaling Technology) and polyclonal antibodies against Y14 (Bethyl), p53 (Proteintech), phospho-histone H3 (Cell Signaling Technology), Cdc20 (Abcam).

**RT-PCR and RT-qPCR**

Total RNA was extracted by using TRIzol reagent (Thermo Fisher Scientific). RNA was treated with RQ1 DNase (Promega) followed by reverse-transcription using SuperScript III kit (Life Technologies). RT-PCR was performed using DreamTaq Green PCR Master Mix (Thermo Fisher Scientific). RT-qPCR was performed using SYBR Green FastMix (Quantabio) in LightCycler480 (Roche). PCR primer sets were listed in Table S3.

**Antibodies**

Immunoblotting was performed as described (Lu et al., 2017) using monoclonal antibodies against cyclin B1 (BD Biosciences), and polyclonal antibodies against cyclin-dependent kinase 1 (CDK1) (Proteintech), p21 (Proteintech) and FLAG (Proteintech).

**Immunofluorescence Staining of Fetal Liver**

Fetal liver on embryonic day 13.5 was isolated, fixed with 4% paraformaldehyde and subjected to perform cryosectioning. The sections were co-stained with FITC-conjugated rat anti-mouse CD41 (BD Biosciences) and polyclonal antibodies against Y14 (Bethyl), p53 (Proteintech) or γH2AX (Novus) overnight at 4˚C, and then were then incubated with appropriate secondary antibodies and stained with Hoechst 33258 (Sigma).

**EdU Labeling**

EdU labeling was performed using Click-iT^TM^ EdU Alexa Fluor^TM^ 647 Flow Cytometry Assay kit (Thermo Fisher Scientific) according to manufacturer’s instructions. In brief, 10 μM EdU was added into the culture medium 1 h before cell harvest. The cells were stained with FITC-conjugated mouse anti-human CD41 (BD Biosciences) and fixed and permeabilized by 4% paraformaldehyde and saponin-based permeabilization/wash reagent for 15 min, respectively. The Click-iT reaction was performed using the Click-iT reaction buffer for 30 min followed by incubation with DAPI and analyzed by flow cytometry as described in the main text. PBS containing 3% BSA was used as the wash buffer between different steps.

**Gene Set Enrichment Analysis**

Gene set enrichment analysis was performed as described (Liberzon et al., 2015) using clusterProfiler (3.10.1). The predefined gene sets were used from Molecular Signatures Database (MSigDB) v6.2 (Subramanian et al., 2005).

**SUPPLEMENTARY TABLES**

**Table S1. Mendelian Ratios of 2-week-old Offspring from Pf4-Cre;*Rbm8a*^f/+^ × *Rbm8a*^f/f^ Cross**

| Genotype | Gender | Expected % | Observed % (N) total=175 |
| --- | --- | --- | --- |
| *Rbm8a*^f/+^ | male | 12.5 | 16.6 (29) |
|  | female | 12.5 | 12.0 (21) |
| *Rbm8a*^f/f^ | male | 12.5 | 10.3 (18) |
|  | female | 12.5 | 13.7 (24) |
| Pf4-Cre;*Rbm8a*^f/+^  (Con) | male | 12.5 | 12.6 (22) |
|  | female | 12.5 | 12.6 (22) |
| Pf4-Cre;*Rbm8a*^f/f^  (*Rbm8a*KO^MK^) | male | 12.5 | 10.3 (18) |
|  | female | 12.5 | 12.0 (21) |

**Table S2. Hematological Analysis of Megakaryocyte-Specific *Rbm8a* knockout mice**

|  | Unit | Con^+^ (N=6) | Con (N=9) | *Rbm8a*KO^MK^ (N=6) |
| --- | --- | --- | --- | --- |
| WBC | 10^3^/µl | 10.68 + 1.66 | 8.95 + 1.15 | 8.54 + 1.78 |
| Neutrophils | 10^3^/µl | 1.63 + 2.18 | 1.25 + 1.47 | 1.92 + 2.49 |
| Lymphocytes | 10^3^/µl | 8.78 + 2.24 | 7.51 + 1.34 | 6.32 + 2.24 |
| Monocytes | 10^3^/µl | 0.05 + 0.03 | 0.04 + 0.01 | 0.04 + 0.02 |
| Eosinophils | 10^3^/µl | 0.22 + 0.8 | 0.15 + 0.03 | 0.25 + 0.10 |
| RBC | 10^6^/µl | 10.46 + 0.31 | 10.56 + 0.35 | 9.98 + 0.54 |
| Platelet | 10^3^/µl | 948.17 + 140.66 | 816.89 + 184.15 | 14.00 + 17.85 ** |
| MPV | fL | 8.08 + 0.19 | 7.88 + 0.24 | 11.45 + 1.20 ** |

Fresh blood was collected in K_2_EDTA tubes and analyzed using an IDEXX ProCyte Dx® Hematology Analyzer. The data represent the mean ± SD of complete blood counts. WBC, white blood cells; RBC, red blood cells; MPV, mean platelet volume (fL, femtolitre); **p < 0.01 (analysis of variance).

**Table S3. Characteristics of *Rbm8a*KO^MK^-p53^Homo^ Mice**

| No. | Gender | PLT (10^3^/μL) | Age of death |
| --- | --- | --- | --- |
| 1 | F | 476 | 4 w |
| 2 | M | - | <3 w |
| 3 | M | - | <3 w |
| 4 | F | 207 | 6 w |
| 5 | F | - | <5 w |
| 6 | F | - | <4 w |
| 7 | M | - | <4 w |
| 8 | F | - | <5 w |
| 9 | F | 596 | 3.5 w |
| 10 | M | - | <3 w |

**Table S4. The List of Primers**

| Experiments/  Gene names | Sequences (5’ to 3’) |
| --- | --- |
| Genotyping | |
| Pf4-Cre-F | CCCATACAGCACACCTTTTG |
| Pf4-Cre-R | TGCACAGTCAGCAGGTT |
| *Rbm8a-*5’ end loxP-F | GAAGATTTCGCCATGGATGAGGATGG |
| *Rbm8a-*5’ end loxP-R | GTTTGTGGATGCTTTCTAGAGTTCCAG |
| *Rbm8a-*3’ end loxP-F | GATCTACCTGCCTCTTCCTCCCA |
| *Rbm8a-*3’ end loxP-R | GACAGATTGAATCCCCCATCCACA |
| *Trp-*5’ end loxP-F | CACAAAAACAGGTTAAACCCAG |
| *Trp-*5’ end loxP-R | AGCACATAGGAGGCAGAGAC |
| *Trp-*3’ end loxP-F | AAGGGGTATGAGGGACAAGG |
| *Trp-*3’ end loxP-R | GAAGACAGAAAAGGGGAGGG |
| RT-qPCR | |
| *ACTB*-F | GCACTCTTGCAGCCTTCCTTCC |
| *ACTB*-R | TGTCACCTTCACCGTTCCAG |
| *ASNS*-F | GGGGCTTGGACTCCAGCTTG |
| *ASNS*-R | GAGCCTGAATGCCTTCCTCA |
| *CD41/ITGA2B*-F | AGGGTGGTGCTGTGTGAG |
| *CD41/ITGA2B*-R | CTGTTCTTGCTCCGTATCTGC |
| *CD61/ITGB3*-F | TGTATGGGACTCAAGATTGGAGAC |
| *CD61/ITGB3*-R | AGCGATGGCTATTAGGTTCAGC |
| *CREB3L3*-F | GGGAGACGAGCTGTGAGC |
| *CREB3L3*-R | TGTCTGAGTGTCGGTTCCTG |
| *CTH*-F | TGGATGATGTGTATGGAGGTACAAACAGG |
| *CTH*-R | GCCTTCAATGTCAATCACCTTCTGGG |
| *CXCL8/IL8*-F | CCTGATTTCTGCAGCTCTGT |
| *CXCL8/IL8*-R | AACTTCTCCACAACCCTCTG |
| *FAM129A*-F | CCAGAACTTCCAGACCACCAA |
| *FAM129A*-R | CGGAATGCAGCGGAAGATT |
| *FcGR1B*-F | GCAAGTGGACACCACAAAGG |
| *FcGR1B*-R | AGTGGCTGTGCCATTGAGAA |
| *FcGR2A*-F | ATCATTGTGGCTGTGGTCATTGC |
| *FcGR2A*-R | TCAGGTAGATGTTTTTATCATCG |
| *FER*-F | TTCGAGGGCACTGGGTTTTC |
| *FER*-R | TTCCCTTGCCCAGTAATTCTCC |
| *HSPA13*-F | ACCGCAGAAGAGTTGGAGGCTGA |
| *HSPA13*-R | TCTGGGGACACTGTGATGGTCTCA |
| *NFATc1*-F | CCCAGATGGCCACCATGT |
| *NFATc1*-R | AGGTCCCGGTCAGTTTTCG |
| *PMAIP1*-F | CAGAGCTGGAAGTCGAGTGT |
| *PMAIP1*-R | AGGAGTCCCCTCATGCAAGT |
| *PRKCD*-F | TTCGGGAAGGTGCTGCTTG |
| *PRKCD*-R | TGCCCTTGCTGTGTAGAAAC |
| *TNFSRF10B*-F | TGCAGTGTCTTTGGTCTCCT |
| *TNFSRF10B*-R | TTCTTGTGAGCTGTGTTGCC |
| *TRIB3*-F | GTCTGGTCCTGCGTGATCTCAA |
| *TRIB3*-R | GTATGAGGCCCGTGAGCTGAGT |

F, forward; R, reverse

**SUPPLEMENTARY FIGURE LEGENDS**

**Figure S1. Knockdown of Y14 Results in Reduced DNA Synthesis. Related to Figure 4.**

HEL cells were transduced with lentivirus encoding control or Y14-specific shRNA followed by mock or PMA treatment as in Figure 4A. Cells were labeled with EdU for 1 h before cell harvest. Megakaryocytes were gated by the CD41 marker. Bar graph shows the percentage of EdU-labeled cells in total CD41^+^ cells or the 2N-4N CD41^+^ cells. Data represent the mean ± SEM.

**Figure S2. *Rbm8a* Knockout Increases the Level of Phosphorylated Histone 3 and γH2AX in Bone Marrow. Related to Figure 4.**

Immunohistochemical staining of phospho-H3 (upper) and γH2AX (lower) in bone-marrow sections of 9-week-old Con and *Rbm8a*KO^MK^ mice. A selected megakaryocyte in the red dashed-line square is magnified in the inset. Scale bars represent 50 μm (insets 20 μm).

**Figure S3. *Rbm8a* Knockout Increases the Level of p53 and γH2AX in Fetal Liver. Related to Figure S2.**

Sections of embryonic day 13.5 fetal liver were double-stained with FITC-conjugated anti-CD41 (green) and an antibody against Y14, p53 or γH2AX (red). DNA was counterstained with Hoechst 33258 (blue). Bar scale, 20 μm.

**Figure S4. p53 Suppresses Y14 mRNA Expression. Related to Figure 6.**

HEK293 cells were transfected with the p53 expression vector for 2 days. Immunoblotting was performed using an antibody against p53, Y14 or GAPDH. RT-PCR was performed using primers listed in Table S4.

**Figure S5. Generation of Megakaryocyte-Specific *Rbm8a* and *Trp53* Double-Knockout Mice. Related to Figure 7.**

(A) Schematic diagram of the strategy to generate megakaryocyte-specific *Rbm8a* and *Trp53* double-knockout mice*.*

(B) PCR conformed the genotypes of *Trp53*^f/+^ and *Trp53*^f/f^-bearing mice using primers as listed in Table S4.

(C) PCR confirmed floxed *Rbm8a* and *Trp53* alleles in bone-marrow cells using 5’ loxP (forward) and 3’ loxP (reverse) primers as listed in Table S4.

(D) Schematic diagrams indicate *loxP* positions and the floxed alleles of *Rbm8a* and *Trp53*. PCR products were subjected to Sanger sequencing analysis.

**Figure S6. Lymph Node Tumors of *Rbm8a*KO^MK^-p53^Homo^. Related to Figure 7.**

A tumor (right) was dissected from the lymph node (left) of *Rbm8a*KO^MK^-p53^Homo^. Bar Scale, 1 cm.

**Figure S7. HE Staining of Bone Marrow Sections from *Trp53*-knockout *Rbm8a*KO^MK^ and Control Mice. Related to Figure 7.**

HE staining of bone marrow sections from *Trp53*^f/+^, *Trp53*^f/f^, *Rbm8a*KO^MK^-p53^Het^ and *Rbm8a*KO^MK^-p53^Homo^ mice. A representative field for each genotype is shown; yellow arrows indicate megakaryocytes (bar scale, 100 µm). Magnified image of the cells in square is shown in Figure 7C.

**Figure S8. Knockdown of Y14 Impairs the Expression of Endomitotic Factors During Megakaryocyte Differentiation. Related to Figure 4.**

(A) Gene Set Enrichment Analysis was performed using data from RNA-seq as in Figure 5A. Normalized enrichment score = –2.48, p < 0.05.

(B) HEL cells were transduced and treated as in Figure 4A. Immunoblotting was performed using antibodies as indicated.

**Figure S9. The p53-M133K Mutant Exhibits Partial Activity in Transcriptional Activation. Related to Figure 6.**

HEL cells were transfected with the empty vector or the p53- or p53 M133K–expressing vector for 2 days. Immunoblotting was performed with an antibody against FLAG, p21, or GAPDH.
