## Supplemental figures for "The Y14-p53 Regulatory Circuit in Megakaryocyte Differentiation and Thrombocytopenia"

### Slide 1
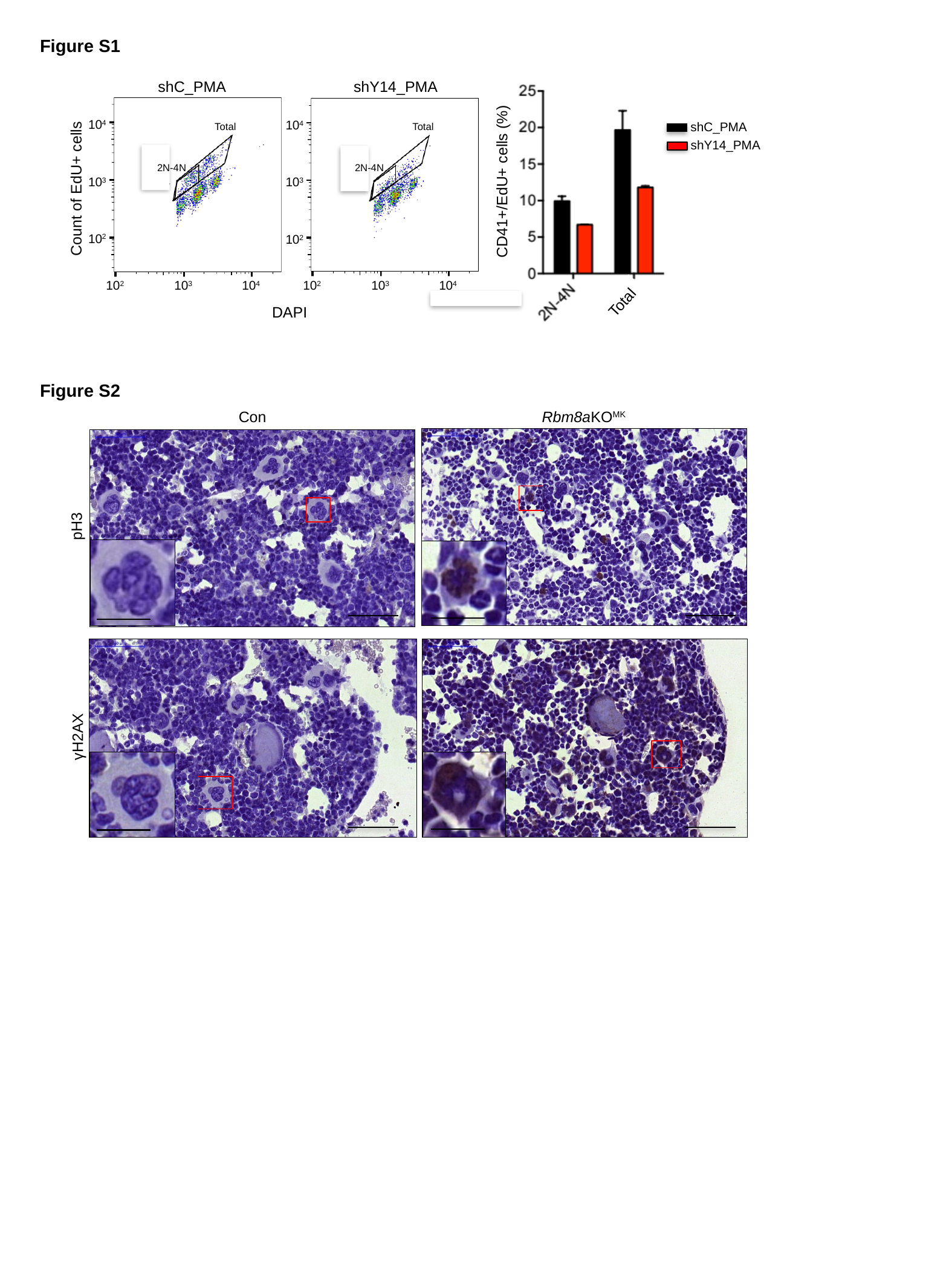

Figure S1
shC_PMA
shY14_PMA
CD41+/EdU+ cells (%)
shC_PMA
shY14_PMA
10^4^
10^4^
Total
Total
2N-4N
2N-4N
10^3^
10^3^
Count of EdU+ cells
10^2^
10^2^
10^2^
10^3^
10^4^
10^2^
10^3^
10^4^
Total
DAPI
Figure S2
Con
Rbm8aKO^MK^
pH3
γH2AX

### Slide 2
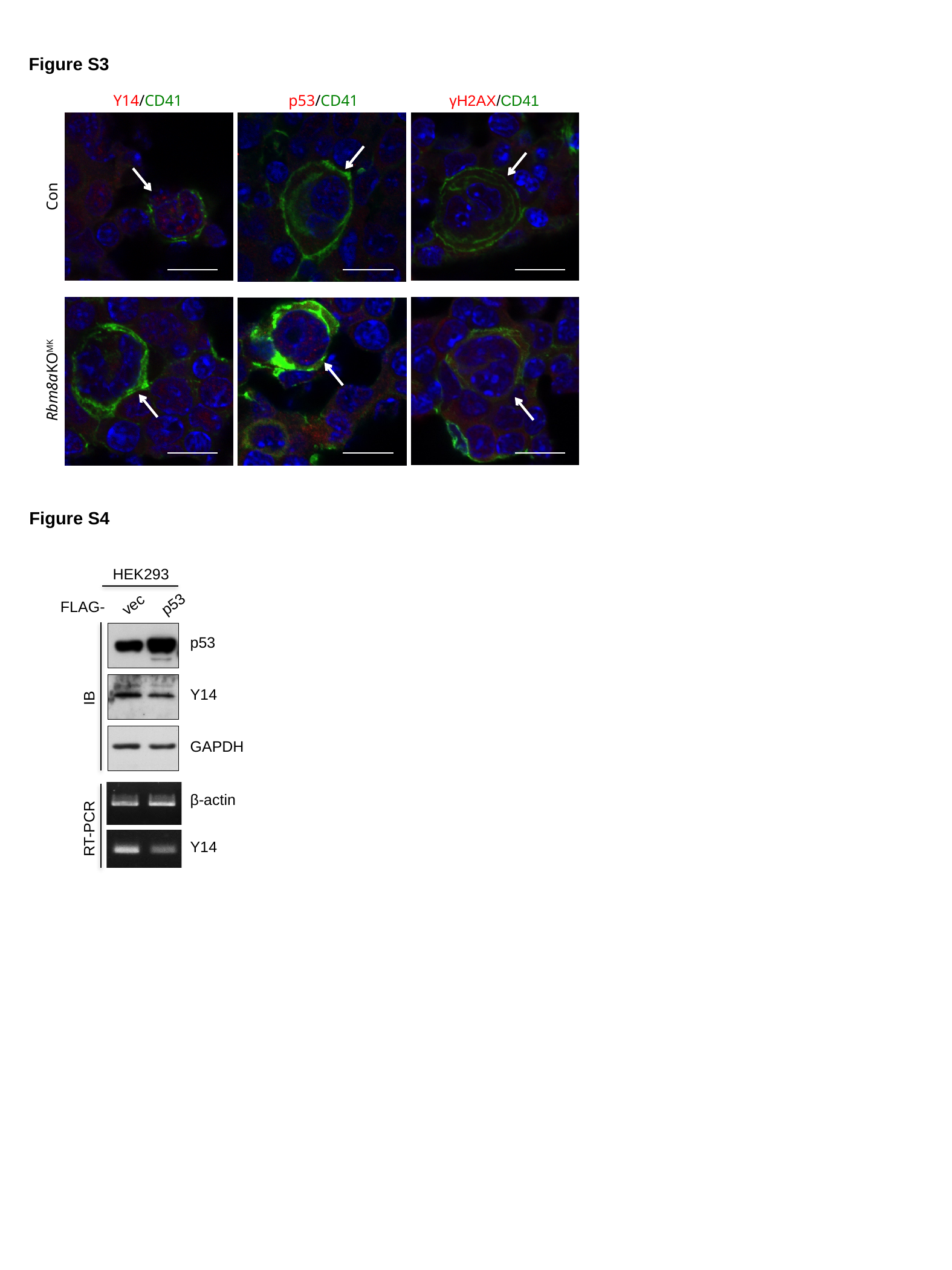

Figure S3
Y14/CD41
p53/CD41
γH2AX/CD41
Con
Rbm8aKO^MK^
Figure S4
HEK293
vec
p53
FLAG-
p53
Y14
IB
GAPDH
β-actin
RT-PCR
Y14

### Slide 3
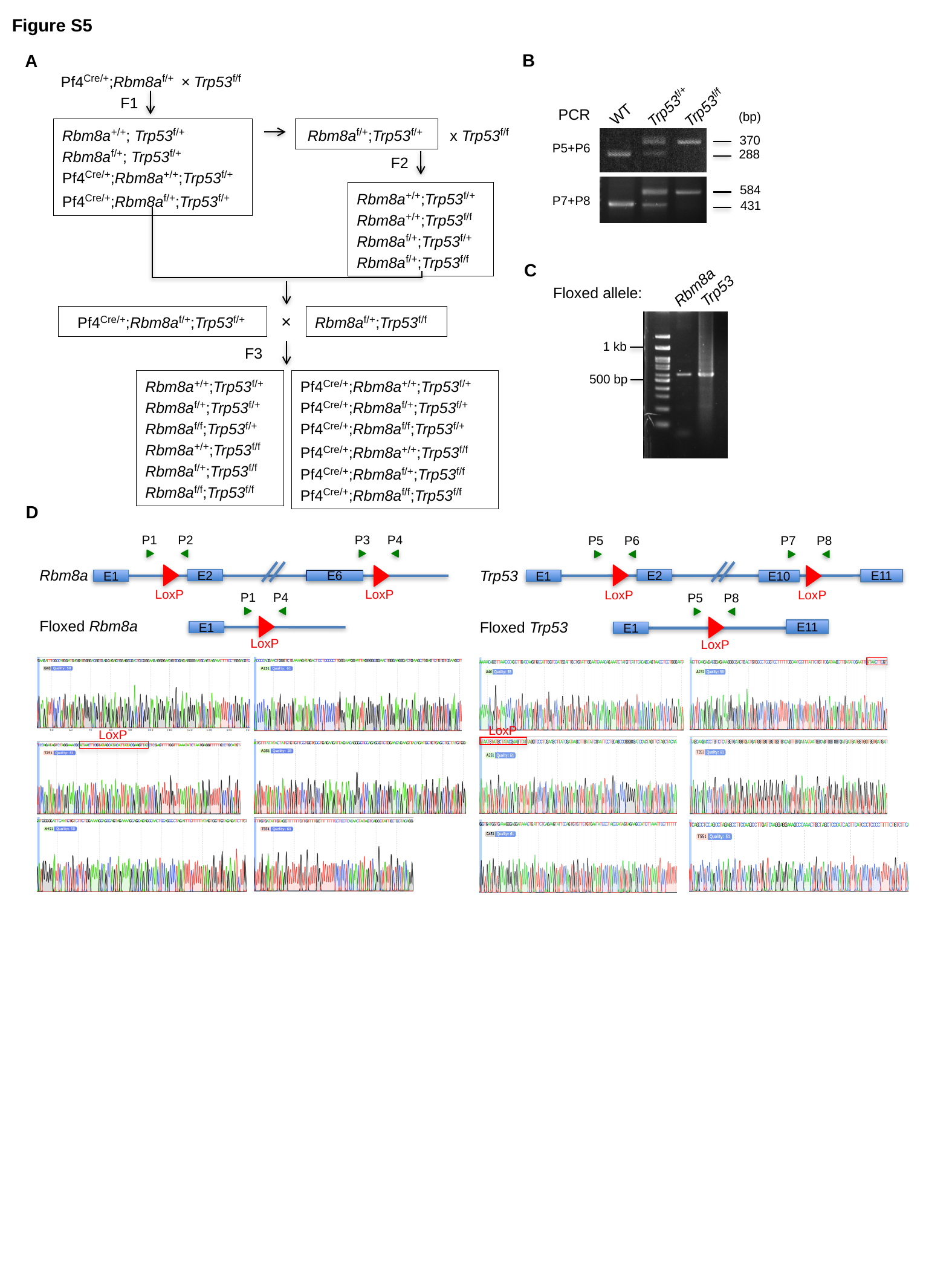

Figure S5
B
A
Pf4^Cre/+^;Rbm8a^f/+^ × Trp53^f/f^
Trp53^f/f^
Trp53^f/+^
WT
PCR
370
P5+P6
288
584
P7+P8
431
F1
(bp)
Rbm8a^+/+^; Trp53^f/+^
Rbm8a^f/+^; Trp53^f/+^
Pf4^Cre/+^;Rbm8a^+/+^;Trp53^f/+^
Pf4^Cre/+^;Rbm8a^f/+^;Trp53^f/+^
Rbm8a^f/+^;Trp53^f/+^
x Trp53^f/f^
F2
Rbm8a^+/+^;Trp53^f/+^
Rbm8a^+/+^;Trp53^f/f^
Rbm8a^f/+^;Trp53^f/+^
Rbm8a^f/+^;Trp53^f/f^
C
Trp53
Rbm8a
1 kb
500 bp
Floxed allele:
Pf4^Cre/+^;Rbm8a^f/+^;Trp53^f/+^
×
Rbm8a^f/+^;Trp53^f/f^
F3
Rbm8a^+/+^;Trp53^f/+^
Rbm8a^f/+^;Trp53^f/+^
Rbm8a^f/f^;Trp53^f/+^
Rbm8a^+/+^;Trp53^f/f^
Rbm8a^f/+^;Trp53^f/f^
Rbm8a^f/f^;Trp53^f/f^
Pf4^Cre/+^;Rbm8a^+/+^;Trp53^f/+^
Pf4^Cre/+^;Rbm8a^f/+^;Trp53^f/+^
Pf4^Cre/+^;Rbm8a^f/f^;Trp53^f/+^
Pf4^Cre/+^;Rbm8a^+/+^;Trp53^f/f^
Pf4^Cre/+^;Rbm8a^f/+^;Trp53^f/f^
Pf4^Cre/+^;Rbm8a^f/f^;Trp53^f/f^
D
P1 P2
P3 P4
Rbm8a
E2
E1
E6
LoxP
LoxP
P1 P4
Floxed Rbm8a
E1
LoxP
LoxP
P5 P6
P7 P8
Trp53
E2
E11
E1
E10
LoxP
LoxP
P5 P8
Floxed Trp53
E11
E1
LoxP
LoxP

### Slide 4
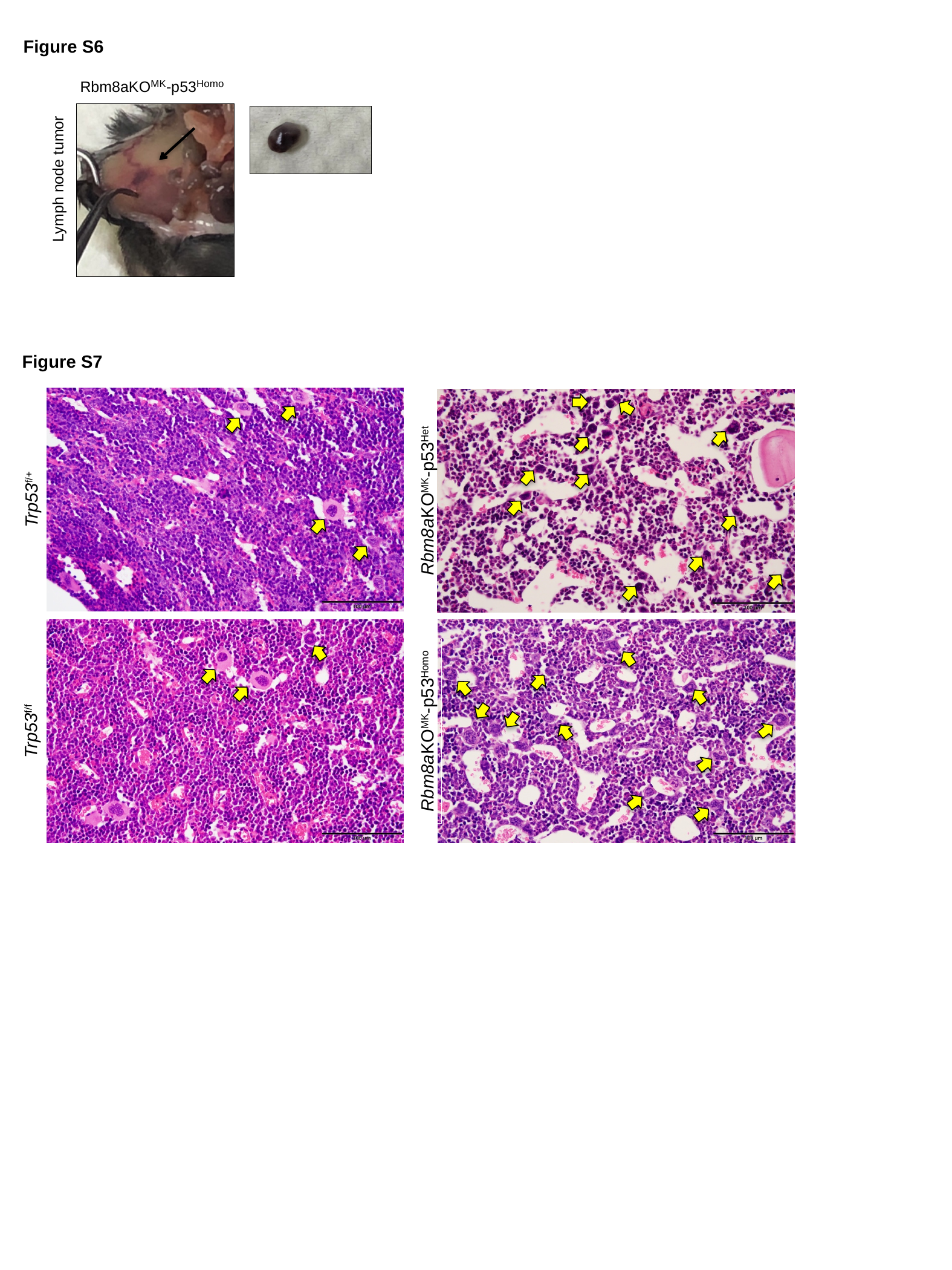

Figure S6
Rbm8aKO^MK^-p53^Homo^
Lymph node tumor
Figure S7
Trp53^f/+^
Rbm8aKO^MK^-p53^Het^
Trp53^f/f^
Rbm8aKO^MK^-p53^Homo^

### Slide 5
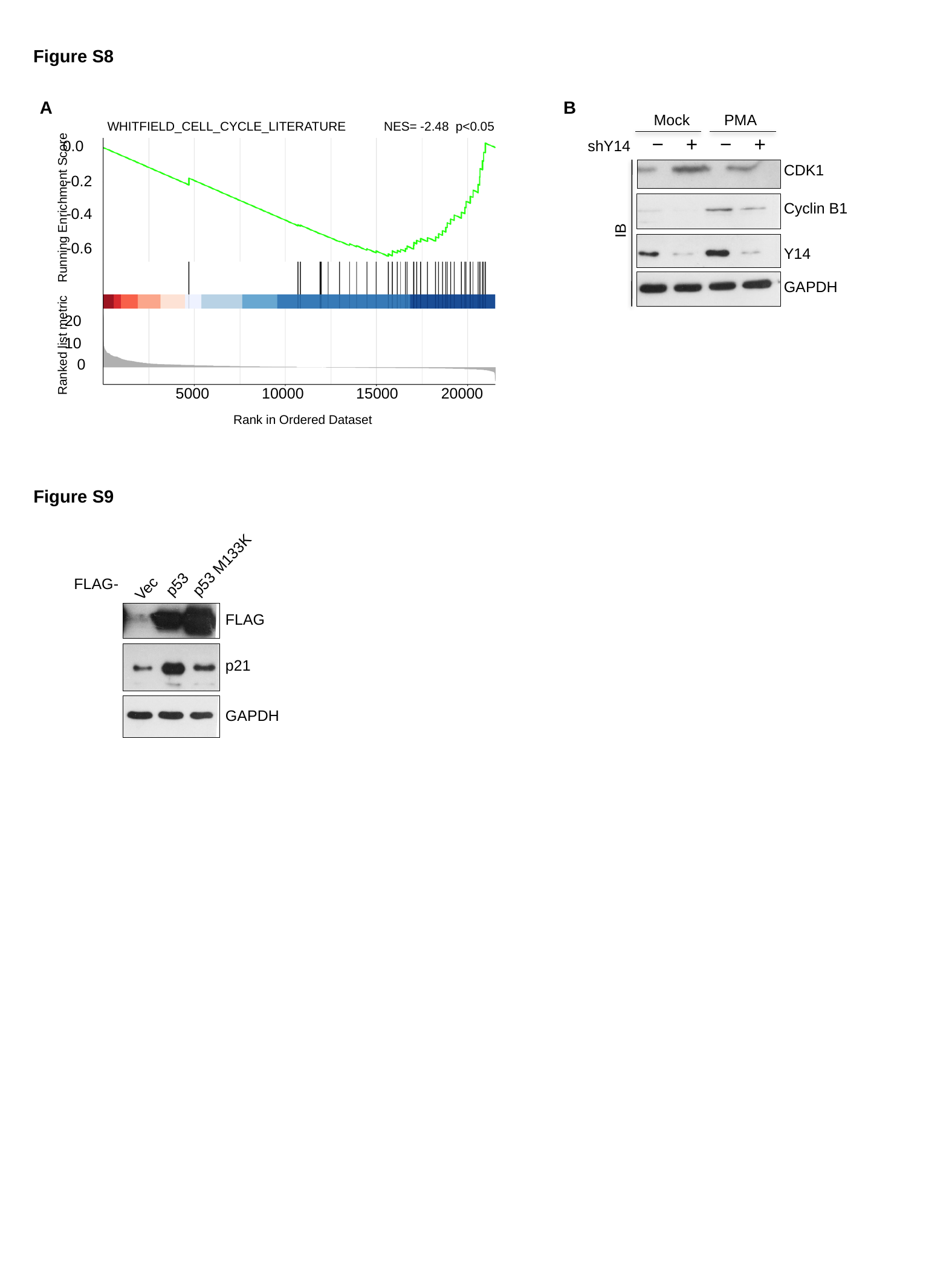

Figure S8
A
B
Mock
PMA
NES= -2.48 p<0.05
0.0
-0.2
-0.4
-0.6
20
10
Ranked list metric
0
5000
10000
15000
20000
Rank in Ordered Dataset
Running Enrichment Score
WHITFIELD_CELL_CYCLE_LITERATURE
shY14 − + − +
CDK1
Cyclin B1
IB
Y14
GAPDH
Figure S9
p53 M133K
p53
Vec
FLAG-
FLAG
p21
GAPDH
